# CryoDRGN-AI: Neural *ab initio* reconstruction of challenging cryo-EM and cryo-ET datasets

**DOI:** 10.1101/2024.05.30.596729

**Authors:** Axel Levy, Rishwanth Raghu, J. Ryan Feathers, Michal Grzadkowski, Frédéric Poitevin, Jake D. Johnston, Francesca Vallese, Oliver Biggs Clarke, Gordon Wetzstein, Ellen D. Zhong

**Affiliations:** Department of Electrical Engineering, Stanford University, Stanford, CA, USA; SLAC National Accelerator Laboratory, Menlo Park, CA, USA; Department of Computer Science, Princeton University, Princeton, NJ, USA; Department of Physiology and Cellular Biophysics, Columbia University, New York, NY, USA; Department of Anesthesiology, Columbia University Irving Medical Center, New York, NY, USA; Irving Institute for Clinical and Translational Research, Columbia University, New York, NY, USA; Structural Biology Initiative, CUNY Advanced Science Research Center, New York, NY, USA

## Abstract

Proteins and other biomolecules form dynamic macromolecular machines that are tightly orchestrated to move, bind, and perform chemistry. Cryo-electron microscopy (cryo-EM) and cryo-electron tomography (cryo-ET) can access the intrinsic heterogeneity of these complexes and are therefore key tools for understanding their function. However, 3D reconstruction of the collected imaging data presents a challenging computational problem, especially without any starting information, a setting termed *ab initio* reconstruction. Here, we introduce cryoDRGN-AI, a method leveraging an expressive neural representation and combining an exhaustive search strategy with gradient-based optimization to process challenging heterogeneous datasets. Using cryoDRGN-AI, we reveal new conformational states in large datasets, reconstruct previously unresolved motions from unfiltered datasets, and demonstrate *ab initio* reconstruction of biomolecular complexes from *in situ* data. With this expressive and scalable model for structure determination, we hope to unlock the full potential of cryo-EM and cryo-ET as a high-throughput tool for structural biology and discovery.

## 1 Introduction

Cryo-electron microscopy (cryo-EM) and cryo-electron tomography (cryo-ET) have transformed structural biology with their ability to visualize biomolecular structures at atomic resolution [1, 2] and uncover the conformations of dynamic molecular machines in solution [3, 4, 5, 6, 7, 8] or directly inside cells [9, 10, 11, 12]. However, the path between cryo-EM data and scientific discoveries is often hindered by complex processing pipelines involving many application-specific algorithms. Automating image processing and making them accessible to nonspecialists can therefore bridge the gap between raw data and scientific knowledge.

Understanding structural variability is essential for elucidating the mechanisms of biomolecular complexes. In cryo-EM and cryo-ET, these complexes, or “particles”, are vitrified from ambient temperatures and thus adopt a range of conformations within the space of accessible low-energy states (*i*.*e*., the conformational space). Heterogeneous reconstruction methods aim to reveal this space from the observed set of noisy and randomly oriented projection images. Recent neural and non-neural algorithms have shown success in the unsupervised reconstruction of structural heterogeneity from cryo-EM images [4, 5, 6, 13, 14] (see **Section 2.7** for further details). Existing tools, however, require preliminary structural information of the imaged complex, either in the form of an initial density map or pre-assigned orientations (*i*.*e*., poses) of the particle images. These initial inputs are usually derived assuming a single underlying structure, which inherently limits the extent of structural variability that can be modeled. For heterogeneous samples, multiple rounds of 3D classification are commonly used to obtain an initial structure that can serve as a reasonable starting point for further refinement. Although 3D classification can operate *ab initio* (*i*.*e*., from a random or uninformative starting point) [15], they assume heterogeneity arises from a small set of independent structures. This simplified model of heterogeneity often does not align with the true distribution of structures and, despite time-consuming iterative processing pipelines, still results in missed conformational states.

In this work, we introduce a deep learning method for *ab initio* reconstruction that can reveal structural variability of dynamic biomolecular complexes from cryo-EM and cryo-ET imaging datasets. Our method, cryoDRGN-AI (Deep Reconstructing Generative Networks - *Ab Initio*), leverages the expressivity of implicit neural representations to model complex distributions of structures from single particle or subtomogram tilt series images, while jointly estimating image poses. To combine speed and robustness to noise, cryoDRGN-AI relies on a new pose estimation strategy that switches from hierarchical grid search to gradient descent. We show that cryoDRGN-AI can reconstruct the conformational and compositional landscape of the assembling bacterial ribosome [16], the precatalytic spliceosome [17], and the SARS CoV-2 spike protein [18] without any prior pose estimation procedures. CryoDRGN-AI is robust to the presence of junk or outlier images and more faithfully resolves the structure of the DSL1/SNARE complex [19] compared to existing processing workflows. Applied on large datasets of highly dynamic molecular machines, cryoDRGN-AI reveals new states of the V-ATPase complex [20] and of the human erythrocyte ankyrin-1 complex [21] that were missed in earlier processing approaches. Finally, with cryoDRGN-AI, we process 2D tilt series collected by cryo-ET and reconstruct three states of the *M. pneumoniae* 70S ribosome during its elongation cycle [22] in a single *ab initio* reconstruction step. CryoDRGN-AI is released as part of the cryoDRGN open-source software package and is available at https://cryodrgnai.cs.princeton.edu/.

## 2 Results

### 2.1 Efficient pose estimation with a two-stage strategy

In single particle cryo-EM and cryo-ET subtomogram averaging, each particle image can be associated with a set of image-specific unknown parameters, called *latent variables* (**Figure 1a**). In the case of *ab initio* heterogeneous reconstruction, this set includes the pose *ϕ*_*i*_ *∈* SO(3) *×* ℝ^2^ – the orientation of the particle with respect to the microscope and 2D translation to compensate for the imperfect centering of the particle in the image – and the latent embedding *z*_*i*_ *∈* ℝ^*d*^ – an abstract low-dimensional vector characterizing the conformational state of the particle. The vector space ℝ^*d*^ – also referred to as the conformational space or latent space – can be used to model both compositional and conformational variation among the set of particles and can thus be seen as a finite union of smooth low-dimensional manifolds. In cryoDRGN-AI, this space is represented by *V*_*θ*_, an element of a parametric family Ω = *{*𝒱_*θ*_, *θ ∈* ℝ^*W*^ *}*. Given a latent embedding *z*_*i*_, 𝒱_*θ*_[*z*_*i*_] : ℝ^3^ *→* ℝrepresents a single electron scattering potential (also called density map). We parameterize the family Ω with a neural network (see **Methods**).

**Figure 1:**
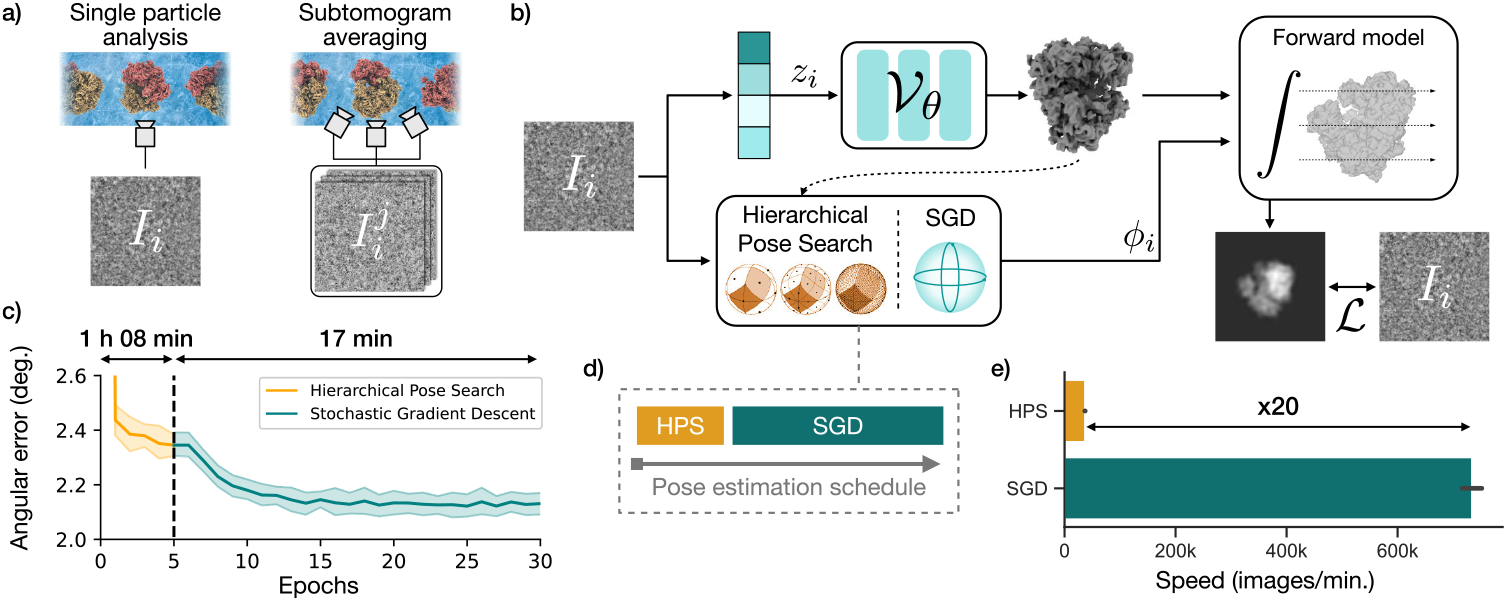
The cryoDRGN-AI method for *ab initio* heterogeneous reconstruction. **a)** CryoDRGN-AI can process single particle images or subtilts for subtomogram averaging. **b)** Architecture overview. Objects in blue are optimized to minimize the loss *ℒ*. For each image, a projection is generated by a differentiable forward model. Poses *ϕ*_*i*_ are first estimated using hierarchical pose search (HPS) and then refined by stochastic gradient descent (SGD). Latent embeddings *z*_*i*_ are optimized by SGD. **c)** Mean out-of-plane angular error during training on a synthetic 80S ribosome dataset (100,000 particles, 128 *×* 128, 3.77 Å/pix, mean *±* std. over 6 runs). **d)** Pose estimation switches to SGD once a set number of images have been processed by HPS. **e)** Gradient descent is 20x faster and leads to more accurate poses since it is not limited by the resolution of the search grid (mean *±* std. over 6 replicates and 5 epochs for HPS, 100 epochs for SGD).

CryoDRGN-AI relies on a new strategy for inferring these latent variables (**Figure 1b**). Instead of using an encoder neural network to map images to latent variables, as in cryoDRGN [5] and other recent works (see **Section 2.7**), we independently store and optimize these variables by gradient descent. The whole pipeline can therefore be seen as an autodecoder [23]. This design choice is motivated by the observation that, in the case of noisy datasets, a neural network tends to *memorize* the images and does not generalize to unseen images [24]. Indeed, in **Supplementary Figure S1**, we show that an encoder trained to predict poses tends to overfit on high-noise datasets (SNR *≤ −*10 dB), preventing true amortization. Thus, instead of using an encoder neural network to memorize latent variable values, we reason that directly storing them in an array is faster, more memory efficient, and more scalable to large datasets.

Pose estimation in cryoDRGN-AI follows a two-stage strategy. First, poses are estimated by a hierarchical pose search algorithm (HPS). During this step, poses are sampled on increasingly fine grids of SO(3) *×* ℝ^2^, projections images are generated through the volume decoder 𝒱_*θ*_[*z*_*i*_] and compared to the observed particle image *I*_*i*_. Although HPS underpins many state-of-the-art methods (see **Section 2.7**), it has two main limitations when adapting to a neural decoder: querying the neural network is computationally expensive, and the final pose accuracy is constrained by the resolution of the finest search grid. To address these challenges, cryoDRGN-AI refines the HPS-estimated poses by stochastic gradient-descent (SGD). We validate this pose estimation strategy on homogeneous reconstruction of a synthetic dataset of 100,000 images of the 80S ribosome (**Methods**). SGD leads to a decrease in the mean angular error (**Figure 1c**), whose effect on the resolution of the final map is quantified in **Supplementary Figure S2**. On this dataset, cryoDRGN-AI pose accuracies and final resolutions are on par with cryoSPARC *ab initio* reconstruction and refinement (**Supplementary Figure S3**). The number of images processed with HPS before switching to SGD is a hyperparameter whose default value is set to max(2*N*, 500, 000) images, where *N* is the number of particle images in the dataset. For images of resolution 128 *×* 128, we find that the processing time is 20 times faster with SGD than with HPS (**Figure 1e**).

### 2.2 Single shot reconstruction on benchmark datasets

To validate cryoDRGN-AI’s *ab initio* heterogeneous reconstruction algorithm, we run cryoDRGN-AI on three experimental benchmark datasets: the pre-catalytic spliceosome (EMPIAR-10180 [17]), the assembling bacterial large ribosomal subunit (EMPIAR-10076 [16]) and the SARS-CoV-2 spike protein [18]. While previous analyses of these datasets typically operated on subsets filtered of junk particles and used consensus poses as input [4, 5, 6, 7, 25, 26, 27, 28, 29], we perform single shot reconstruction, *i*.*e*., a single round of training, on the deposited particle stack, fully *ab initio*.

In **Figure 2**, a visualization of the latent embeddings inferred by cryoDRGN-AI (reduced to 2D via Principal Component Analysis (PCA) or Uniform Manifold Approximation and Projection (UMAP) [30]) is shown on the left, and representative density maps are shown on the right. On the EMPIAR-10180 dataset [17], cryoDRGN-AI reconstructs the continuous bending motion of the pre-catalytic spliceosome while clustering a set of particle images corresponding to broken or denatured density maps in the latent space (**Figure 2a, S4**). CryoDRGN-AI recovers the compositional heterogeneity of the assembly states of the bacterial large ribosomal subunit in the EMPIAR-10076 [16] dataset, while also grouping previously unclassified and discarded particles in the latent space (**Figure 2b**). Finally, on a much smaller protein complex of the SARS-CoV-2 spike protein (<500 kDa) [18], we show that cryoDRGN-AI can reconstruct both the closed and open states of the spike protein fully *ab initio* and resolve the conformational change of the receptor binding domain (**Figure 2c**). In **Supplementary Figure S5**, we provide additional results on a synthetic dataset with strong conformational heterogeneity for which cryoDRGN-AI provides a better reconstruction than cryoSPARC’s multiclass *ab initio* workflow.

**Figure 2:**
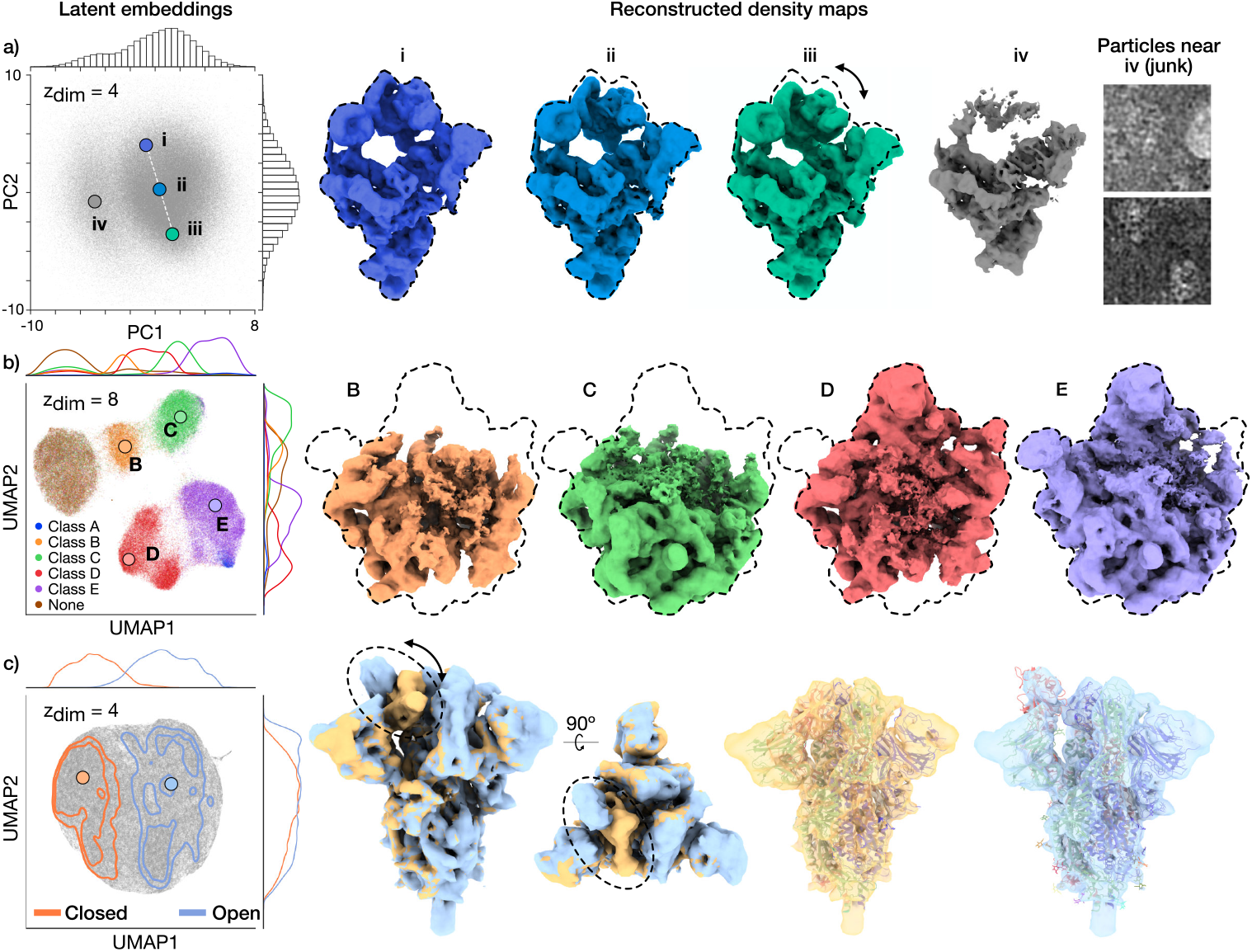
CryoDRGN-AI *ab initio* heterogeneous reconstruction of unfiltered benchmark datasets. **a)** Latent embeddings visualized with PCA and reconstructed maps for the precatalytic spliceosome dataset (EMPIAR-10180 [17], 2.6 MDa, 128 *×* 128, 4.25 Å/pix., 327,490 particles). Dashed lines indicate outlines of the extended spliceosome. **b)** UMAP visualization of latent embeddings and reconstructed maps for the assembling bacterial large ribosomal sub-unit dataset (EMPIAR-10076 [16], approx. 2.1 to 3.3 MDa, 256 *×* 256, 1.64 Å/pix., 131,899 particles). Latent embeddings are colored by their published labels. Dashed lines indicate outlines of the fully mature 50S ribosome. **c)** UMAP visualization of latent embeddings and reconstructed maps for the SARS-CoV-2 spike protein dataset [18] (438 kDa, 128 *×* 128, 3.28 Å/pix., 369,429 particles). Reconstructed density maps of the closed and open states of the receptor binding domain with docked atomic models (PDB:6VXX, PDB:6VYB).

### 2.3 CryoDRGN-AI is robust to junk particles

We next sought to characterize cryoDRGN-AI on a dataset where a majority of particles are outliers that do not contribute to the final structure. We run cryoDRGN-AI on a dataset of the DSL1/SNARE complex (EMPIAR-11846) [19], a 255-kDa complex involved in the membrane fusion reaction of eukaryotic cells. In DAmico *et al*., a total of 214,511 particles were obtained after particle picking and an initial filtering via 2D classification. An iterative reconstruction and classification workflow exploring subsets of the dataset eventually yielded a low-resolution density map. This initial map then seeded several rounds of 3D classification, resulting in a final 6.2 Å consensus reconstruction from 49,947 particles.

We run cryoDRGN-AI on this dataset and observe a distribution of density maps containing the fully intact complex as well as broken particles. We classify the images among three groups by *k*-means clustering in the latent space (**Figure 3a**). By visual inspection, we select 75,854 particles from the cluster associated with the most resolved structure and run a second *ab initio* reconstruction on this subset of particles. After this second round, reconstructed density maps reveal a continuous hinging motion of the DSL1/SNARE complex (**Figure 3b**). For comparison, we show in **Supplementary Figure S6** that simple cryoSPARC-based workflows (multiclass *ab initio* and 2D classification) fail to reconstruct the full structural variability of the complex.

**Figure 3:**
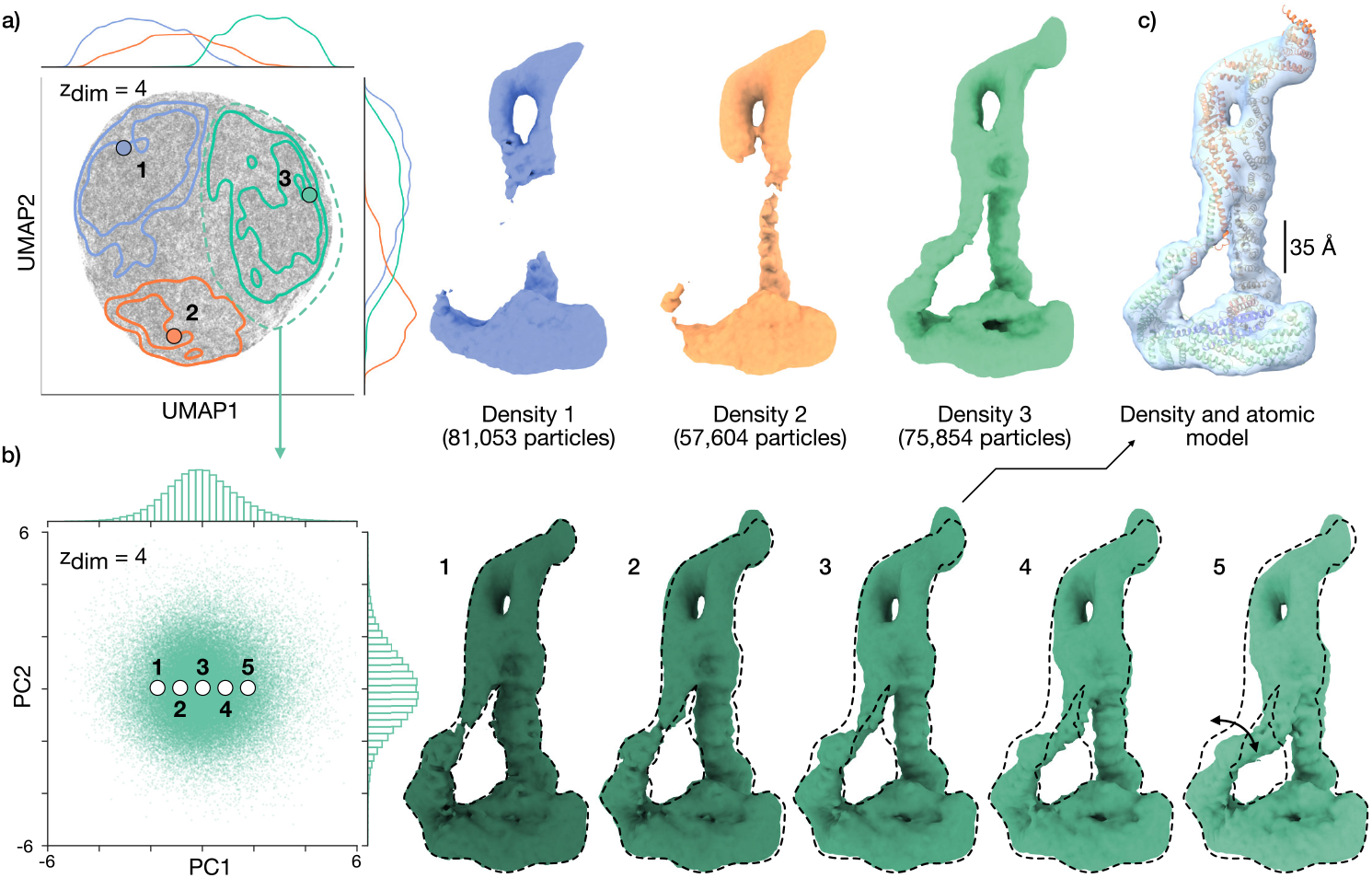
CryoDRGN-AI *ab initio* reconstruction of the DSL1/SNARE complex [19] dataset containing majority junk particles. **a)** UMAP visualization of latent embeddings and reconstructed maps for the full dataset (214,511 particles, 128 *×* 128, 3.47 Å/pix.) with cryoDRGN-AI. The latent embeddings are clustered with *k*-means (*k* = 3). Particles associated with the green cluster are selected. **b)** CryoDRGN-AI latent embeddings visualized with PCA and reconstructed maps for 75,854 selected particles from **a**. Additional density maps are shown in **Supplementary Video 1. c)** Density map from **b** with docked atomic model (PDB: 8EKI).

### 2.4 CryoDRGN-AI captures diverse sources of 3D variability

We next evaluate cryoDRGN-AI on a dataset containing a large degree of compositional and conformational heterogeneity, as well as junk particles: the porcine kidney V-ATPase complex (EMPIAR-10874 [20]), a transmembrane proton pump involved in multiple signaling pathways. Through initial particle picking and 2D classification on the deposited micrographs, 267,216 particles were obtained.

Running cryoDRGN-AI, we observe a distinct cluster in the latent space corresponding to broken particles (**Figure 4a, d**). We filter out these particles using cryoDRGN-AI’s interactive lasso tool and train a new *ab initio* heterogeneous model on the remaining 177,481 particles. Sampled density maps show three conformational states of the complex, characterized by different orientations of the central rotor domains (**Figure 4b, e**). CryoDRGN-AI also resolves compositional heterogeneity in rotary state 2, showing that the signaling protein mEAK-7 can bind to the front of the complex. Additionally, we reveal a previously undescribed state corresponding to mEAK-7 bound to the back of rotary state 3. We observe that the latent space clusters by the binding location (or non-binding) of mEAK-7 (**Figure 4c**). The unbound cluster can be further decomposed into three clusters, associated with the three rotary states (**Supplementary Figure S7**). We validate each state with a voxel-based backprojection of the selected particles and compute a half-map Fourier shell correlation (**Supplementary Figure S7**).

**Figure 4:**
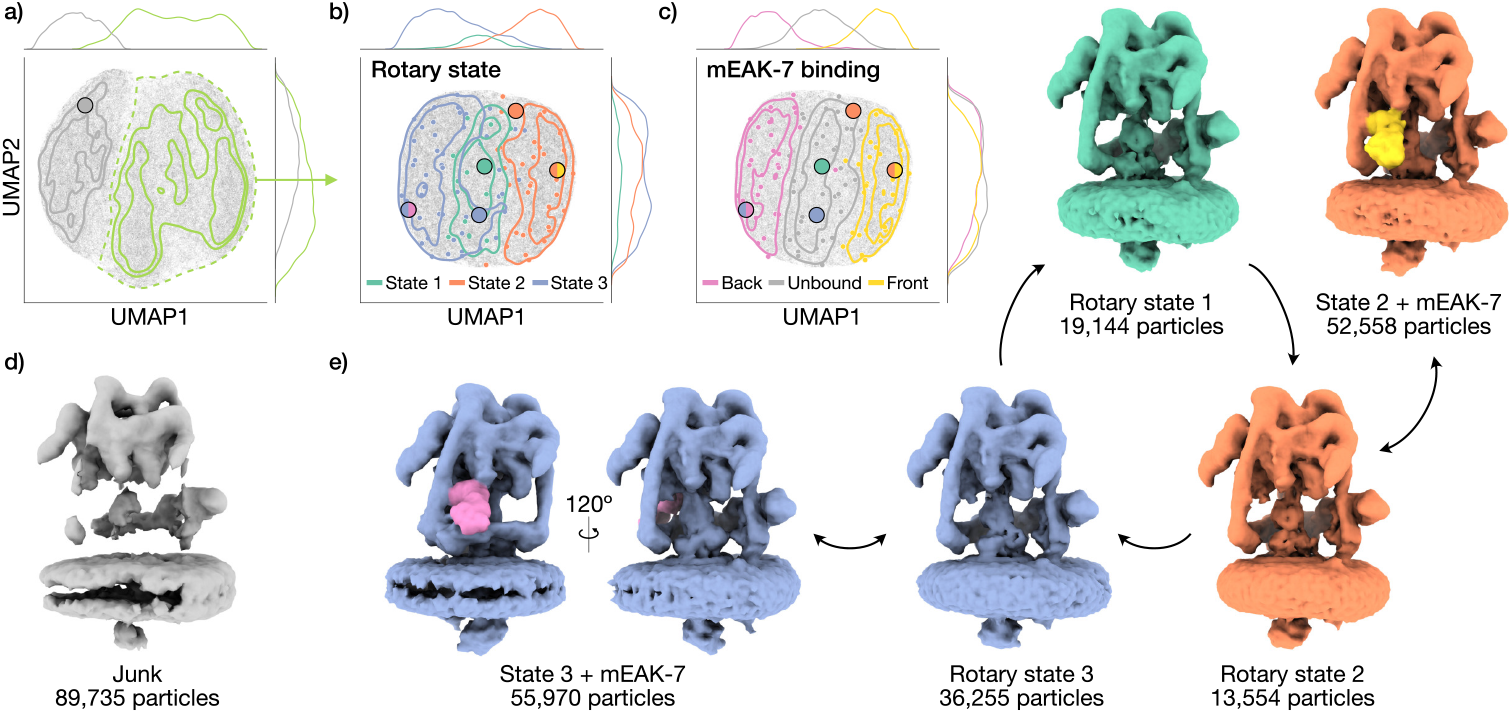
CryoDRGN-AI *ab initio* reconstruction of the V-ATPase complex. **a)** UMAP visualization of cryoDRGN-AI latent embeddings of the full dataset (EMPIAR-10874 [20], 267,216 particles, 128 *×* 128, 3.97 Å/pix.). Selected particles are in green. **b, c)** UMAP visualization of cryoDRGN-AI latent embeddings on the filtered dataset (177,481 particles) with three rotary states (**b**) or mEAK-7 binding location (**c**). Gaussian kernel density estimates are overlaid along with visually classified *k*-means centroids (*k* = 100) as small circles, and points corresponding to the maps in panel (**e**) as large circles. **d)** Sampled density map showing a broken complex from the unfiltered dataset. **e)** Sampled density maps of the three rotary states and mEAK-7 binding.

### 2.5 New state of the human erythrocyte ankyrin-1 complex

An important aspect of cryoDRGN-AI is its ability to process large cryo-EM datasets in an *ab initio* setting where rare states appear in sufficient quantity for detection and 3D reconstruction. We process a dataset of the human erythrocyte ankyrin-1 complex (EMPIAR-11043 [21]), a large membrane-embedded complex important for the shape and stability of red blood cells. The deposited dataset contains more than 700,000 picked particles and was shown to contain six different conformationally and compositionally varying states.

**Figure 5** shows the results of *ab initio* heterogeneous reconstruction with cryoDRGN-AI on the full dataset. Different forms of the complex can be identified by sampling latent embeddings. We run *k*-means clustering (*k* = 6) on the latent embeddings. By evaluating *V*_*θ*_ on the *k*-means centroids, we reconstruct the six previously described classes [21] (**Figure 5a**). Additional conformational heterogeneity can be visualized in the same trained cryoDRGN-AI model by sampling a series of latent embeddings linearly interpolating the populated area between points 2a and 2b, corresponding to a rotation of the ankyrin with respect to the micelle (**Figure 5b**). Increasing the number of sampled density maps to 100 (via *k*-means clustering) revealed a structure of the “supercomplex” state (**Figure 5a**), which was hypothesized but not shown in Vallese *et al*. [21]. This rare state simultaneously contains the Rhesus heterotrimer, aquaporin-1, the three band 3 I-II-III dimers, and an unknown protein Y, that could correspond to the unknown protein X revealed in class 4 by Vallese *et al*. We validated the presence of this new structure by running an independent homogeneous reconstruction on a subset of 20,000 particles (**Figure 5c** and **Methods**).

**Figure 5:**
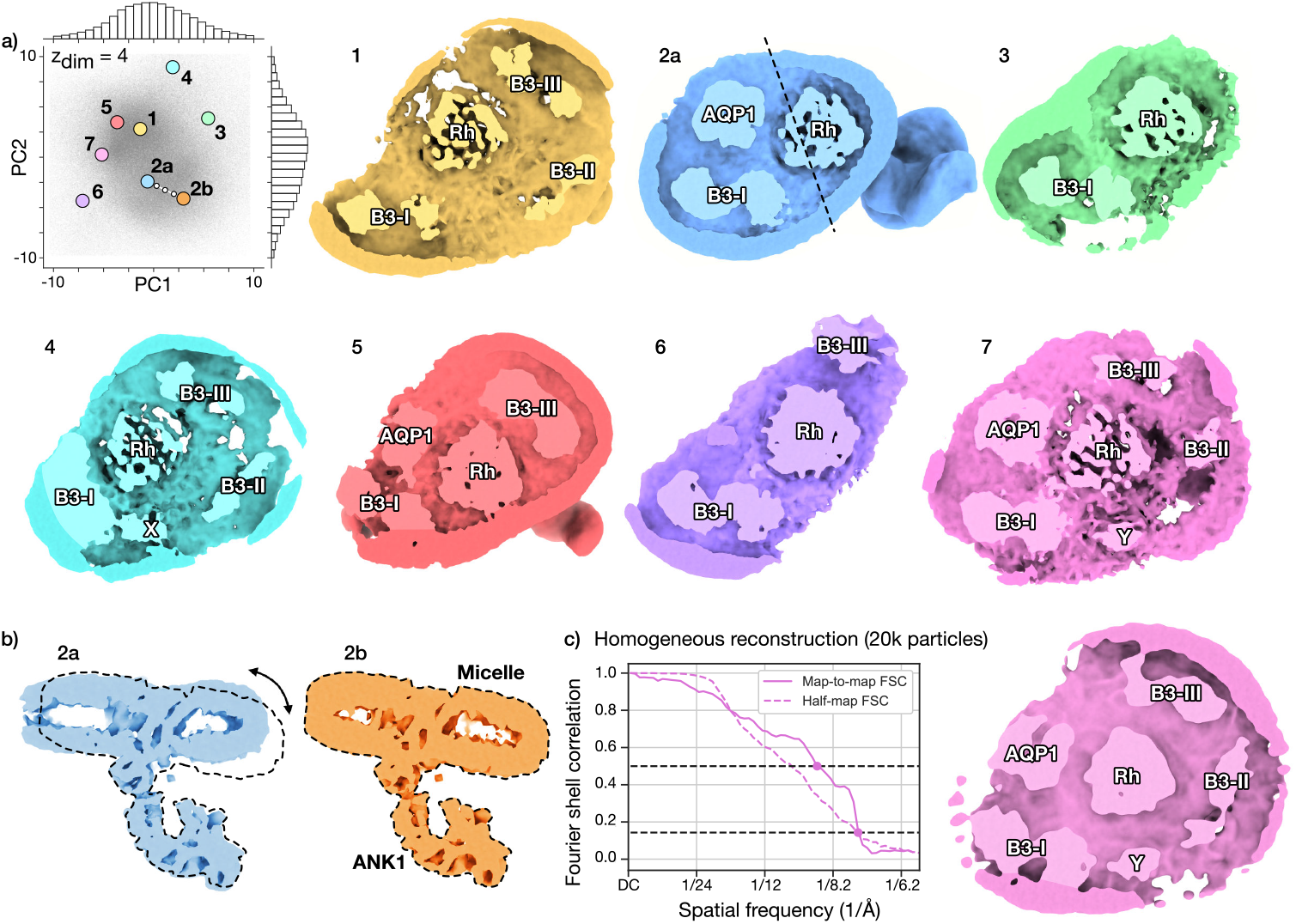
CryoDRGN-AI *ab initio* reconstruction of the human erythrocyte ankyrin-1 complex and identification of a new “supercomplex” state. **a)** CryoDRGN-AI latent embeddings visualized with PCA and reconstructed maps for the full dataset (EMPIAR-11043 [21], 710,437 particles, 128 *×* 128, 2.92 Å/pix.). Reconstructed density maps correspond to the six published classes [21] (1-6), which are distinguished by micelle composition. CryoDRGN-AI also reveals the presence of a new “supercomplex” state (7), which simultaneously contains the rhesus heterotrimer (Rh), the aquaporin (AQP1), the band 3-I (B3-I), the band 3-II (B3-II), the band 3-III (B3-III) dimers and an unknown protein Y in the micelle. Additional density maps shown in **Supplementary Video 2. b)** Linear trajectory in the populated region between 2a and 2b reveals a continuous rotation of the ankyrin (ANK1) relative to the micelle. Plane of view shown as dotted line in 2a in panel **a. c)** Validation of the supercomplex structure. A homogeneous reconstruction in cryoSPARC [15] on 20k particles with poses from cryoDRGN-AI. Map-to-map and half-map FSC curves.

Recent versions of the cryoSPARC software allow 3D classification with large numbers of classes (*e*.*g*., an order of magnitude greater than previous standard settings). As cryoDRGN-AI estimates poses, we performed 3D classification with cryoDRGN-AI poses and reproduced the supercomplex in one of 80 classes (**Supplementary Figure S8**), providing additional validation for this new structure.

### 2.6 CryoDRGN-AI for *ab initio* subtomogram averaging

Finally, we apply cryoDRGN-AI to *ab initio* subtomogram averaging (STA) of tilt series images obtained from cryo-ET and recover the structural and spatial variability of the 70S ribosome *in situ*. In cryoDRGN-AI’s *ab initio* STA, the relative orientations between subtilt images are constrained by the tilting scheme during hierarchical pose search but this constraint is relaxed when switching to stochastic gradient descent to compensate for potential jitter and sample deformation during the imaging process.

**Figure 6** shows the results of an *ab initio* heterogeneous subtomogram averaging on a dataset of the *Mycoplasma pneumoniae* 70S ribosome (EMPIAR-10499 [22]), using 11 tilts per particle. We analyze the influence of the number of tilts on pose error in **Supplementary Figure S9**. By visual inspection of 100 density maps sampled by *k*-means clustering, we reconstruct three known representative states of the ribosome during the translation elongation cycle: the “P state” with a tRNA in the P site, the “EF-Tu, P state” with a tRNA attached to an elongation factor and the “A, P state” where the tRNA has moved to the A site (**Figure 6a, b**). We subsequently run a voxel-based backprojection with the particles associated with the “A, P state” and show the resulting density maps in **Supplementary Figure S9** and **Supplementary Video 3**. We show in **Supplementary Figure S10** that the set of “junk” particles is consistently classified as outliers in replicas of the experiment. The distribution of the translation states is visualized within a representative tomogram in **Figure 6c**. Additionally, cryoDRGN-AI resolves heterogeneity in the cellular context of each particle, for example producing density maps containing neighboring ribosomes in polysomes (**Figure 6d**). Overall, these results highlight the breadth of cryoDRGN-AI’s expressive deep learning-based heterogeneity reconstruction algorithm for *ab initio* reconstruction across a number of challenging cryo-EM and cryo-ET datasets.

**Figure 6:**
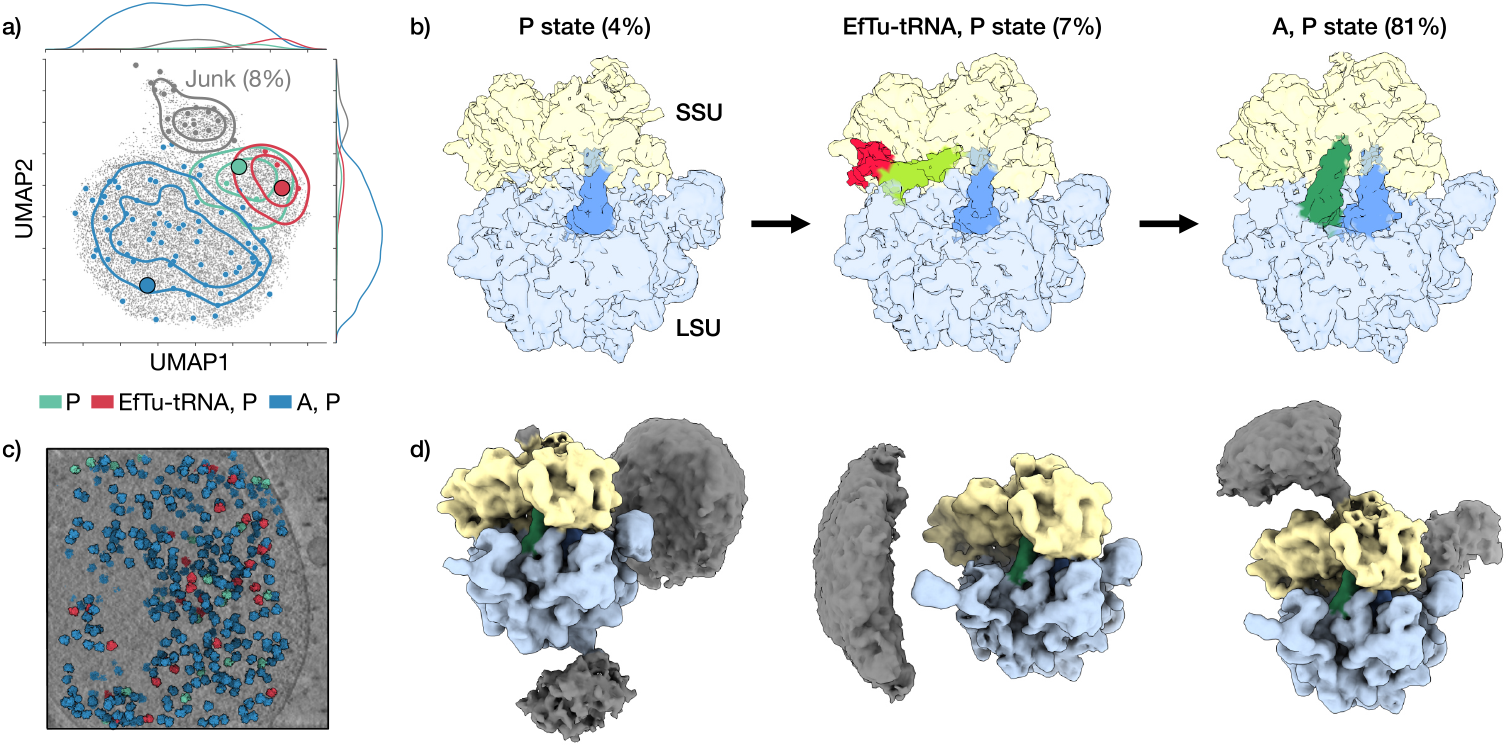
Single shot heterogeneous *ab initio* subtomogram averaging with cryoDRGN-AI. **a)** UMAP visualization of cryoDRGN-AI latent embeddings of a dataset of the *M. pneumoniae* 70S ribosome (EMPIAR-10499 [22]). The 100 maps obtained by *k*-means clustering are classified by visual inspection according to the content of the tRNA sites, between the small sub-unit (SSU) and the large sub-unit (LSU). On top of a 2D scatterplot of the latent embeddings, the classified centroids and a Gaussian KDE of the distribution of states are shown. Large circles indicate the latent embeddings of maps shown in panel **b. b)** Three intermediate states of the ribosome during the translation elongation cycle. **c)** Visualization of a representative *M. pneumoniae* tomogram with ribosomes colored by their class assignments from panel **a. d)** Sampled maps displaying density for neighboring ribosomes in polysomes.

### 2.7 Related Work

#### Modeling heterogeneity in cryo-EM

Recent methods for heterogeneous cryo-EM reconstruction have explored the possibility of using linear [4, 26] and nonlinear [3, 31, 32, 33, 34, 35] mappings between latents and 3D structures. Nonlinear methods can be further categorized depending on the support they use for representing density maps: neural networks [5, 14], 3D Zernike polynomials [28], Gaussian mixture models [6, 36] or flow fields [13]. Normal mode-based approaches [37, 38, 39, 40] introduce a linear and physically interpretable mapping between latents and atomic coordinates. However, these methods require either a coarse initialization of the density map or the poses to be provided by a pre-processing step of homogeneous reconstruction. In practice, this sequential approach can be error-prone due to the presence of junk particles or strong compositional heterogeneity preventing alignment to a single reference structure.

#### *Ab initio* single particle analysis

While early algorithms for heterogeneous reconstruction were based on expectation-maximization [41, 42, 43] and thus required careful initialization for accurate convergence, stochastic gradient-based optimization techniques were later introduced for *ab initio* reconstruction and released as part of the cryoSPARC software [15, 44]. These methods, termed “multiclass *ab initio*” or “3D classification with alignments”, approximate the conformational landscape with a small set of independent voxel grids while inferring poses with a search-based procedure. While these methods can be used in practice to recover diverse structures from complex samples [45, 46], they are fundamentally limited to modeling discrete heterogeneity over a user-defined number of classes. An early version of cryoDRGN [14], cryoDRGN2 [27] and cryoFIRE [47] introduced preliminary neural *ab initio* methods for heterogeneous reconstruction. CryoDRGN2 incorporates a hierarchical pose search strategy [27], but was ultimately not evaluated on its ability to scale to large datasets. CryoFIRE uses an autoencoder to estimate poses but amortized inference is not well-suited for highly noisy datasets (*e*.*g*., for smaller complexes), where the encoder may memorize the data [24] (**Supplementary Figure S1**). We propose a two-step pose estimation strategy and an autodecoding architecture to explicitly address the limitations of previous neural approaches, and in doing so, reveal both the compositional and conformational heterogeneity of large, unfiltered datasets. We note that recently-published works [48] also showed the benefits of using a two-step optimization strategy for pose estimation, starting with amortized inference. While these works support the design of our approach, they have only addressed the problem of homogeneous reconstruction.

#### Subtomogram averaging

Subtomogram averaging traditionally operates on “3D subtomograms”, *i*.*e*., sub-volumes extracted from the 3D tomogram that is obtained from a voxel-based backprojection of 2D tilt series images [38, 49, 50, 51, 52]. As samples cannot be imaged at high tilt angles, a “missing wedge” in Fourier space leads to degraded high-frequency information in the 3D subtomograms. Recently, methods using per-particle and per-tilt corrections [12, 53, 54, 55, 56] have been developed as an alternative approach for subtomogram averaging. In particular, cryoDRGN-ET [12] extends cryoDRGN to infer conformational landscapes directly from particle tilt images but requires known poses. CryoDRGN-AI expands this capability further to *ab initio* STA, in hopes of enabling faster and more streamlined processing of electron tomography data.

## 3 Discussion

In this work, we demonstrate new capabilities for recovering 3D structure and variability from diverse, challenging single-particle cryo-EM and cryo-ET subtomogram datasets. Unlike most existing methods for heterogeneity analysis, cryoDRGN-AI can be trained in an *ab initio* setting, while leveraging an expressive implicit neural representation to reconstruct compositionally and conformationally diverse structures. We find that our hybrid pose search and autodecoding algorithm leads to successful optimization of the model in challenging settings, in particular in the regime of large, unfiltered cryo-EM datasets. For example, we identified a new state of the human erythrocyte ankyrin-1 complex (**Figure 5**) and of the V-ATPase complex (**Figure 4**) missed in prior processing approaches. We additionally recovered the structure and motion of the DSL1/SNARE complex from a dataset containing a significant fraction of junk particles (**Figure 3**), a common occurrence in cryo-EM datasets that typically leads to complex and *ad hoc* processing pipelines [57]. Furthermore, we show that simpler sequential workflows with traditional tools fail to recover the full variability of the complex, demonstrating the potential utility of incorporating *ab initio* heterogeneous reconstruction in an earlier stage of image processing (**Supplementary Figure S6**). Finally, cryoDRGN-AI provides the capability for joint inference of poses and heterogeneity in cryo-ET STA (**Figure 6**), enabling reconstruction of datasets with complex heterogeneity inherent to the cellular milieu.

How should the learned distribution of structures be interpreted and analyzed? Although we do not use any explicit regularization scheme as in variational approaches to generative modeling [58], we empirically observe that similar density maps tend to be close to each other in latent space. As such, we found that compositional heterogeneity could be revealed from clusters in the learned latent embeddings and conformational heterogeneity could be visualized through continuous trajectories in the latent space (**Figure 2**), following the same analysis pipeline as cryoDRGN [5, 59]. We note that the layout of the cryoDRGN-AI latent space can differ from that produced by cryoDRGN [5] due to its latent variable optimization via an autodecoder framework. We refer to Jeon *et al*. [60] for a survey and comparison of heterogeneous reconstruction methods. As in all deep learning approaches that learn an abstract latent space for heterogeneity, the distribution of embeddings does not have a direct physical meaning (*e*.*g*., as a conformational energy landscape), and imbuing a thermodynamic interpretation to distances in latent space remains an open challenge.

Over the past decade, cryo-EM has become a dominant approach for experimental biomolecular structure determination. While recent machine learning breakthroughs have transformed our ability to computationally predict protein structure from sequence [61, 62] and design protein structures *de novo* [63, 64], *these tools propose structures purely in silico*, and therefore require experimental validation for many downstream applications or scientific study. With cryoDRGN-AI, we seek to contribute to automated, reproducible pipelines that remove the manual, subjective decisions in cryo-EM data processing, thus increasing the throughput of experimental structure determination. In addition, we hope to expand the capabilities of cryo-EM reconstruction to more complex samples where the current paradigm of assigning poses separately before heterogeneity analysis is insufficient. Lastly, we anticipate that these new capabilities in biomolecular structure determination will drive the creation of larger and more information-rich training datasets for other AI methods in structural biology and provide major advances in our understanding and engineering of biological molecules.

## Acknowledgements

We thank Mike Dunne, Ramya Rangan, and members of the Zhong lab for assistance on the manuscript. We thank Kevin DAmico and Fred Hughson for guidance on structure interpretation. We thank Bridget Carragher, Clint Potter, and members of the Chan Zuckerberg Imaging Institute for feedback on the cryo-ET experiments. We thank Vineet Bansal for software engineering support. We acknowledge the use of computational resources from the SLAC Shared Scientific Data Facility and from Princeton Research Computing, which is consortium of groups led by the Princeton Institute for Computational Science and Engineering (PICSciE) and Office of Information Technology’s Research Computing. Some of this work was performed at the Simons Electron Microscopy Center at the New York Structural Biology Center, with support from the Simons Foundation (SF349247), NIH (U24 GM129539), and NY State Assembly. A.L. and F.P. were supported by the Department of Energy, Laboratory Directed Research and Development program at SLAC National Accelerator Laboratory, under contract DE-AC02-76SF00515. O.B.C was supported by the National Heart, Lung, and Blood Institute (NHLBI) of the NIH through grant 1R01HL168178. The Zhong lab gratefully acknowledges support from the Princeton Catalysis Initiative, Princeton School of Engineering and Applied Sciences, and Generate:Biomedicines. The funders had no role in study design, data collection and analysis, decision to publish or preparation of the manuscript.

## Author Contributions

A.L., F.P., G.W., and E.Z. conceived of the work; A.L., R.R., J.R.F. and E.Z implemented the methods and performed the computational experiments; A.L., R.R., M.G., and E.Z. developed the software package; J.J. preprocessed subtomogram data; F.V. and O.B.C. analyzed the results on the ankyrin dataset; A.L., R.R. and E.Z. wrote the manuscript with feedback from all the authors; G.W. and E.Z. supervised the project.

## Methods

### Image formation model

In single-particle cryo-EM, an image *I*_*i*_ is considered to be a random realization following

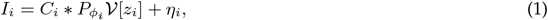

where *C*_*i*_ is the (discretized) Contrast Transfer Function (CTF), *\** the convolution operator and *η*_*i*_ additive white Gaussian noise [65]. *V* maps conformations (*z ∈* ℝ^*d*^) to 3D density maps (*V* [*z*] : ℝ^3^ *→*ℝ). The operator *P*_*ϕ*_ represents an orthographic projection of the map 𝒱 [*z*_*i*_] from the pose *ϕ* = (*R*, **t**) *∈* SO(3) *×* ℝ^2^, followed with a discretization of the projected image. For *V* : ℝ^3^ *→*ℝ,

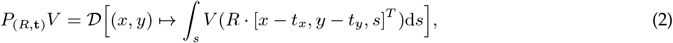

where, for *I* : ℝ^2^ *→* ℝand a *D*^2^ grid *{*(*x*_*m*_, *y*_*n*_)*}*_*m,n∈{*1,…,*D}*_,

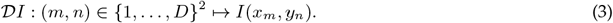

In order to avoid the computation of integrals, we simulate the image formation model in Hartley space, thereby allowing the use of the Fourier Slice Theorem. In Hartley space, the above image formation model can be re-written

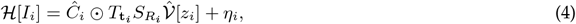

where ℋ is the discrete Hartley transform, Ĉ_*i*_ is the Hartley transform of *C*_*i*_, *⊙* the element-wise multiplication, 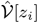 the Hartley transform of 𝒱 [*z*_*i*_] and *S*_*R*_ corresponds to a “slicing” operation,

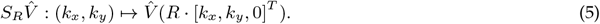

*T*_**t**_ corresponds to a translation in Hartley space. Given *H* : ℝ^2^ *→* ℝand a *D*^2^ grid 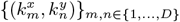,it is defined by

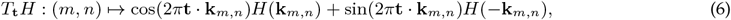

where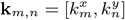. In practice, we use a regular grid centered around the origin, *i*.*e*.,

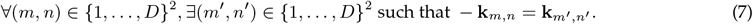

### Volume representation

In cryoDRGN-AI, the ensemble of density maps 𝒱_*θ*_ : ℝ^*d×*3^ *→* ℝis represented by a neural network with parameters *θ*. Conditioned on a latent embedding *z*_*i*_ *∈* ℝ^*d*^, *V*_*θ*_[*z*_*i*_] : ℝ^3^ *→* ℝrepresents the Hartley transform of the 3D electron scattering potential of a single particle. The frequency coordinate **k** *∈* [*−*0.5, 0.5]^3^ is expanded in a sinusoidal basis using Fourier features [66]. We use 64 base frequencies randomly sampled from a 3D Gaussian distribution of standard deviation 0.5. The neural network follows the same architecture as in cryoDRGN [5]: the concatenation of *z*_*i*_ with the positionally-encoded frequency is passed through 3 hidden residual layers [67] of size 256 with ReLU non-linearities. The last layer is linear and provides a one-dimensional output.

### Training and optimization

CryoDRGN-AI reproduces the image formation model in Hartley space and aims at minimizing the reconstruction error in Hartley space (negative log-likelihood of the observations):

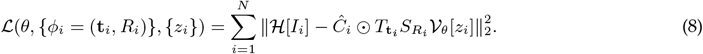

In cryoDRGN-AI, 𝒱_*θ*_ : ℝ^*d×*3^ *→* ℝis represented by a neural network, the poses *ϕ*_*i*_ are optimized with a two-step strategy and the latent embeddings *z*_*i*_ are independently optimized by gradient descent.

The weights of the neural network are randomly initialized with the default initialization scheme in PyTorch [68], and latent embeddings are sampled from a *d*-dimensional Gaussian distribution of standard deviation 0.1. Optimization starts with a “pretraining phase” during which the weights of the neural network are optimized by gradient descent on the objective function, using fixed random poses (uniform over SO(3)) and latent embeddings for 10,000 images by default. We use the Adam optimizer [69] with a learning rate of 0.0001 and a batch size of 32. After the pretraining phase, the latent embeddings are then optimized using the Adam optimizer and a learning rate of 0.01, while the poses are estimated using hierarchical pose search (HPS). To compensate for the high memory requirements of HPS, the batch size is reduced to 8. Once max(2*N*, 5 *×* 10^5^) images have been processed, where *N* is the number of images in the dataset, the estimated poses are refined with stochastic gradient descent (SGD). Poses are optimized using the Adam optimizer with a learning rate of 0.001 and the batch size is increased to 256. All experiments were run on 4 A100s NVIDIA GPUs using data parallelization.

### Hierarchical pose search

The hierarchical pose search (HPS) formulation is adapted from Zhong *et al*. [27]. The reprojection error is defined as

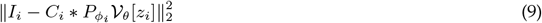

and is directly computed in Hartley space. The first search step is done on a predefined uniform grid over SO(3) *×* ℝ^2^ (4,608 rotations with 15 degrees spacing and 49 translations on a 7 *×* 7 grid in [*−*10 pix., 10 pix.]). The top 8 rotations minimizing the reprojection error are kept and refined with a local search over 8 neighboring rotations at half the grid resolution. The extent of the translation grid is halved, but remains on a 7 *×* 7 grid. Hierarchical search proceeds for 4 additional steps. The images are band-limited during pose search and the cutoff frequency increases linearly from *k*_min_ to *k*_max_ (*k*_min_ = 6, *k*_max_ = 16 in (image length)^*−*1^). Grids are parameterized using the Hopf fibration [70], product of the Healpix [71] grid on the 2-sphere and a regular grid on the circle.

### Additional details for subtomogram averaging

Similar to **Equation 1**, the image formation model for subtilt image 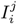 corresponding to the *j*-th tilt of the *i*-th particle follows:

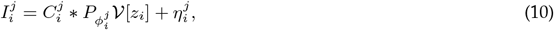

where the CTFs, poses and noises are tilt-dependent while the latent embeddings are only particle-dependent.

To account for accumulated radiation damage in tomography, we expand the CTF model for 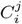 to account for lower signal-to-noise ratio (SNR) in tilts collected at later time-points and at higher tilt angles. We include a dose exposure correction to account for frequency-dependent signal attenuation in later tilt images, as described in [72]. Additionally, since sample thickness effectively increases at higher tilt angles leading to decreasing SNR for these tilts, we further multiply the CTF by the cosine of the tilt angle [73].

For pose search in subtomogram averaging, the reprojection error (**Equation 9**) is replaced with

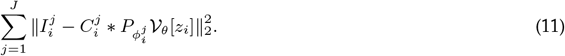

In this case, a single pose is optimized per particle, and the known tilting scheme constrains the relative orientations between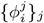. This constraint is relaxed during SGD phase, and poses are optimized independently in order to compensate for potential jitter and sample deformation during the imaging process. We use the first 11 tilt images from the dose-symmetric tilting scheme, balancing accuracy (**Supplementary Figure S9**) with the increase in compute cost for pose search of additional tilt images.

For subtomogram averaging, max(2*N*, 1.5 *×* 10^5^) particles are processed during HPS, the learning rate for poses is 0.00001 during SGD, and the batch size during SGD is reduced to 32. Additionally, while processing the first 50,000 particles during HPS, conformations are randomly sampled from a *d*-dimensional Gaussian distribution with standard deviation 0.1 at each epoch.

### Analysis of the conformational landscape

We follow the ‘cryodrgn analyze’ pipeline to analyze the distribution of reconstructed volumes, where the predicted latent embeddings are visually inspected in 2D using PCA or UMAP [30] and density maps are sampled at *k*-means cluster centers of the latent embeddings [5]. We generally find it easier to interpret continuous deformations with PCA (as in **Figure 2a**) and reveal compositional heterogeneity with UMAP (as in **Figure 2b, c**). Density maps are visualized with UCSF ChimeraX [74] and rendered at a single isosurface level of the potential in all figures.

No measures are explicitly taken to disentangle poses and latent embeddings (*i*.*e*., to guarantee that a motion in latent space does not represent only a rigid transformation of the density map). We refer to Klindt *et al*. [75] for an in-depth discussion on ways to evaluate and potentially correct pose-conformation entanglement.

## Metrics

### Per-image FSC

For synthetic datasets, where ground truth density maps are available for each particle image, we quantify the accuracy of a heterogeneous reconstruction using the “per-image FSC” [14]. It corresponds to the Fourier Shell Correlation between the ground truth density map and the predicted 3D map associated with each image, averaged across the dataset. In cryoDRGN-AI, the predicted map is obtained by feeding the latent embedding of a given image to the decoder. In cryoSPARC, each image is classified among a finite number of possible classes and each class is associated with a predicted density map.

### Pose accuracy

To compute rotation accuracy, the estimated rotations **r**_*i*_ are first aligned to the reference rotations **R**_*i*_ using 100 candidates 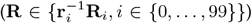 and keeping the minimizer of the mean Frobenius norm of the difference between the reference rotation matrices and the estimated aligned rotation matrices **r**_*i*_**R**. The rotation matrices are then decomposed into viewing directions and viewing angles using the Euler decomposition in the canonical reference frame. The “out-of-plane” error is then defined as the angular error between the estimated (aligned) viewing directions and the reference viewing directions. The “in-plane” error is defined as the angular error between the viewing angles.

When computing 2D translation errors, we account for the fact that the origin of the 3D reference frame is unknown. Given a set of estimated rotation matrices **r**_*i*_ *∈* ℝ^3*×*3^, a set of estimated 2D translations **t**_*i*_ *∈* ℝ^2^ and a set of reference 2D translations **T**_*i*_ *∈* ℝ^2^, the “optimal 3D shift” **u** *∈* ℝ^3^ is defined as the minimizer of the mean square error of the corrected translations

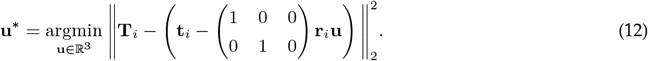

The minimizer is obtained with a numpy least-square solver and the corrected translations are defined as

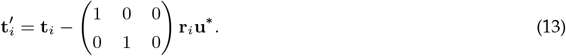

### Validation in cryoSPARC

We used cryoSPARC version 4.4 [15] to validate the structure of the new “supercomplex” state in the ankyrin-1 complex dataset. We computed the L2 distance between the 710,437 reconstructed maps and kept the images associated with the 20,000 maps that were the closest from the identified “supercomplex” state (in L2 distance). Validation was obtained running a homogeneous reconstruction job in cryoSPARC, based on the poses predicted by cryoDRGN-AI. We report two FSC curves: half-map and map-to-map (*i*.*e*., we compare cryoSPARC’s density map to cryoDRGN-AI’s density map). We run cryoSPARC 3D classification using 80 classes, with poses provided by cryoDRGN-AI, at a target resolution of 12 Å, for 40 O-EM epochs, with initial low pass filtering to 60 Å, class similarity of 0.1, auto-tuning of initial class similarity as True, and all other parameters set to their default values. The resulting density map of the supercomplex was identified by visual inspection of classes. A comparison to cryoDRGN-AI’s latent embeddings is provided in **Supplementary Figure S8**.

### Analysis of pose encoder

To evaluate the capability of a neural-based encoder to predict latent variables (here, poses) purely from images (**Fig. S1**), a neural network mapping particle images to poses is supervised on synthetic cryo-EM datasets of the 80S ribosome with different noise levels. Each dataset is normalized in pixel space and split into a training set (20k particles) and a test set (10k particles). The neural network is built with a stack of 2D convolutional layers (kernel size size of 3, “reflection” mode for padding) interlaced with ReLU activation functions at every layer and a group normalization layer (32 groups) followed by an average pooling layer every other layer. The sequential CHW sizes are: 1 *×* 128 *×* 128, 32 *×* 128 *×* 128, 32 *×* 64 *×* 64, 64 *×* 64 *×* 64, 64 *×* 32 *×* 32, 128 *×* 32 *×* 32, 128 *×* 16 *×* 16, 256 *×* 16 *×* 16, 256 *×* 8 *×* 8, 512 *×* 8 *×* 8. The last layer is a tanh activation function followed by a linear layer with 6 output dimensions. The output is interpreted as a rotation using the *S*^2^ *× S*^2^ parameterization [76] and converted into a quaternion. The neural network is optimized with the Adam optimizer, a learning rate of 0.0001, and a batch size of 32 for 250 epochs. The loss for a pair of poses (**q, q**^*′*^), represented as quaternions, is defined by

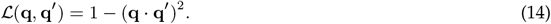

### Data

A summary of all the datasets analyzed in this study and cryoDRGN-AI dataset-specific training settings are found in **Table 1**. We also provide a summary of the parameters of the deposited particle stacks in **Table 2**, for the datasets that were processed from micrographs.

**Table 1:** Datasets and cryoDRGN-AI training settings. *D* refers to the box size in pixels.

| Dataset | EMPIAR | SPA/ET | $D$ | Å/pix. | Particles | Tilts | $z_{\text{dim}}$ | Epochs | Time | GPUs |
| --- | --- | --- | --- | --- | --- | --- | --- | --- | --- | --- |
| Pre-catalytic spliceosome [17] | 10180 | SPA | 128 | 4.25 | 327,490 | – | 4 | 100 | 04:41 h | 4xA100 |
| Assembling ribosome [16] | 10076 | SPA | 256 | 1.64 | 131,899 | – | 8 | 104 | 11:36 h | 4xA100 |
| Spike protein [18] | N/A | SPA | 128 | 3.28 | 369,429 | – | 4 | 100 | 05:19 h | 4xA100 |
| DSL1/SNARE (run 1) [19] | 11846 | SPA | 128 | 3.47 | 214,511 | – | 4 | 100 | 03:43 h | 4xA100 |
| DSL1/SNARE (run 2) [19] | 11846 | SPA | 128 | 3.47 | 75,854 | – | 4 | 100 | 02:11 h | 4xA100 |
| V-ATPase (run 1) [20] | 10874 | SPA | 128 | 3.97 | 267,216 | – | 4 | 101 | 08:40 h | 4xA100 |
| V-ATPase (run 2) [20] | 10874 | SPA | 128 | 3.97 | 177,481 | – | 4 | 102 | 07:05 h | 4xA100 |
| Ankyrin [21] | 11043 | SPA | 128 | 2.92 | 710,437 | – | 4 | 720 | 48:04 h | 4xA100 |
| 70S Ribosome [22] | 10499 | ET | 128 | 3.90 | 18,466 | 11 | 16 | 108 | 46:02 h | 4xA100 |
| Synthetic 80S ribosome [77] | N/A | SPA | 128 | 3.77 | 100,000 | – | – | 30 | 01:25 h | 4xA100 |
| Synthetic 1D rotation [5] | N/A | SPA | 128 | 6.00 | 50,000 | – | 8 | 100 | 03:02 h | 4xA100 |

**Table 2:**
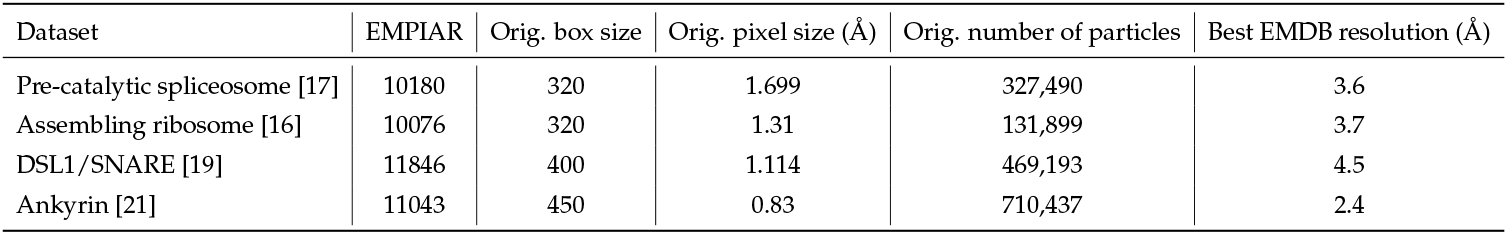
Original parameters of the datasets processed from picked particles deposited on EMPIAR.

| Dataset | EMPIAR | Orig. box size | Orig. pixel size (Å) | Orig. number of particles | Best EMDB resolution (Å) |
| --- | --- | --- | --- | --- | --- |
| Pre-catalytic spliceosome [17] | 10180 | 320 | 1.699 | 327,490 | 3.6 |
| Assembling ribosome [16] | 10076 | 320 | 1.31 | 131,899 | 3.7 |
| DSL1/SNARE [19] | 11846 | 400 | 1.114 | 469,193 | 4.5 |
| Ankyrin [21] | 11043 | 450 | 0.83 | 710,437 | 2.4 |

### Synthetic 80S ribosome

We generate synthetic datasets of the 80S ribosome using the same procedure as Levy *et al*. [77] by following the image formation model (**Equation 1**). The electron scattering potential was simulated in ChimeraX [74] at 6.0 Å resolution from a combination of two atomic models: the small subunit (PDB:3J7A) and the large subunit (PDB:3J79) [78]. The dataset contains 100,000 particles with an image size of 128 *×* 128 and a pixel size of 3.77 Å. Rotations are uniformly sampled over SO(3) and images are centered (no translations). CTF defocus values are randomly sampled using log-normal distributions following Levy *et al*. [77]. Zero-mean white Gaussian noise is added with a standard deviation of 0.5. Example images are shown in **Supplementary Figure S3**.

### Synthetic 1D rotation

We used the synthetic dataset introduced in Zhong *et al*.[5] to simulate strong conformational heterogeneity. The dataset contains 50,000 particles. 50 density maps were generated along a 1D reaction coordinate defined by rotating a dihedral angle in the atomic model of a hypothetical protein complex (0.03 rad. increment, **Supplementary Figure S5a**). Density maps were generated using the molmap command in ChimeraX [74] at a grid spacing of 6 Å per pixel, resolution of 12 Å and a box size of 128^3^ voxels. Each density map is projected 1,000 times using random poses (translations of *±*10 pixels). CTF and noise at a signal-to-noise ratio of 0.1 were added to the dataset. The CTF defocus values were randomly sampled from EMPIAR-10028 [78].

### Pre-catalytic spliceosome

The EMPIAR-10180 [17] dataset was downsampled to 128 *×* 128 from the original image size of 320 *×* 320, giving a pixel size of 4.25 Å/pix. We process the full dataset of deposited particles (327,490 particles).

### Assembling 50S ribosome

The EMPIAR-10076 [16] dataset was downsampled to 256 *×* 256 from the original image size of 320 *×* 320, giving a pixel size of 1.64 Å/pix. We process the full dataset of deposited particles (131,899 particles).

### SARS-CoV-2 spike protein

The dataset from Walls *et al*. [18] was provided courtesy of the authors. Particle images were downsampled to 128 *×* 128 from the original resolution 400 *×* 400, giving a pixel size of 3.28 Å/pix. We process the full dataset of 369,429 particles.

### DSL1/SNARE complex

The EMPIAR-11846 [19] dataset was downsampled to 128 *×* 128 from the original image size of 400 *×* 400, giving a pixel size of 3.47 Å/pix. The original dataset contains 286,801 picked particles and was filtered down to 214,511 particles by visual inspection of the latent space obtained with cryoDRGN [5].

### V-ATPase complex

V-ATPase particles were obtained from reanalysis of EMPIAR-10874 [20] dataset entry #10 (unaligned multiframe movies of Pig Kidney V-ATPase bound to mEAK-7 with ATP collected using Titan Krios and Falcon4). Initial preprocessing prior to cryoDRGN-AI analysis was performed in RELION5 [79]. Briefly, 5,498 movies were aligned and motion corrected using the RELION implementation of MotionCor2. 1,517,258 particles were picked using 2D class average templates generated from classification of an initial Laplacian-of-Gaussian blob pick on a subset of images. Iterative 2D classification was used to remove classes that appeared to contain false positive picks and empty micelles resulting in a particle stack containing 267,216 particle images downsampled to 128 *×* 128 (3.97 Å/pix).

### Ankyrin-1 complex

The EMPIAR-11043 [21] dataset was downsampled to 128 *×* 128 from the original image size of 450 *×* 450, giving a pixel size of 2.92 Å/pix. We process the full dataset of ankyrin complex particles (710,437 particles). For this dataset, we extended the training time to 48 hours to maximize the chances of detecting any novel states (see **Table 1**).

### *Mycoplasma pneumoniae* 70S ribosome

Subtomograms were picked and processed from EMPIAR-10499 [22] with the same protocol as performed in Rangan *et al*. [12]. The dataset contains 18,466 particles with 41 tilts each (757,106 subtilt images). We keep the first 11 tilts (3 degrees between consecutive tilts, total coverage of *±*15 degrees) at a resolution of 128 *×* 128 (3.9 Å/pix.). The voxel-based backprojection of the selected “A, P” state particles in **Supplementary Figure S9** was performed at a resolution of 294 *×* 294 (1.7 Å/pix.) and low pass filtered in ChimeraX [74] using an isotropic Gaussian kernel of standard deviation 1.5 Å.

## Supplementary Tables

## Supplementary Figures

**Supplementary Figure S1:**
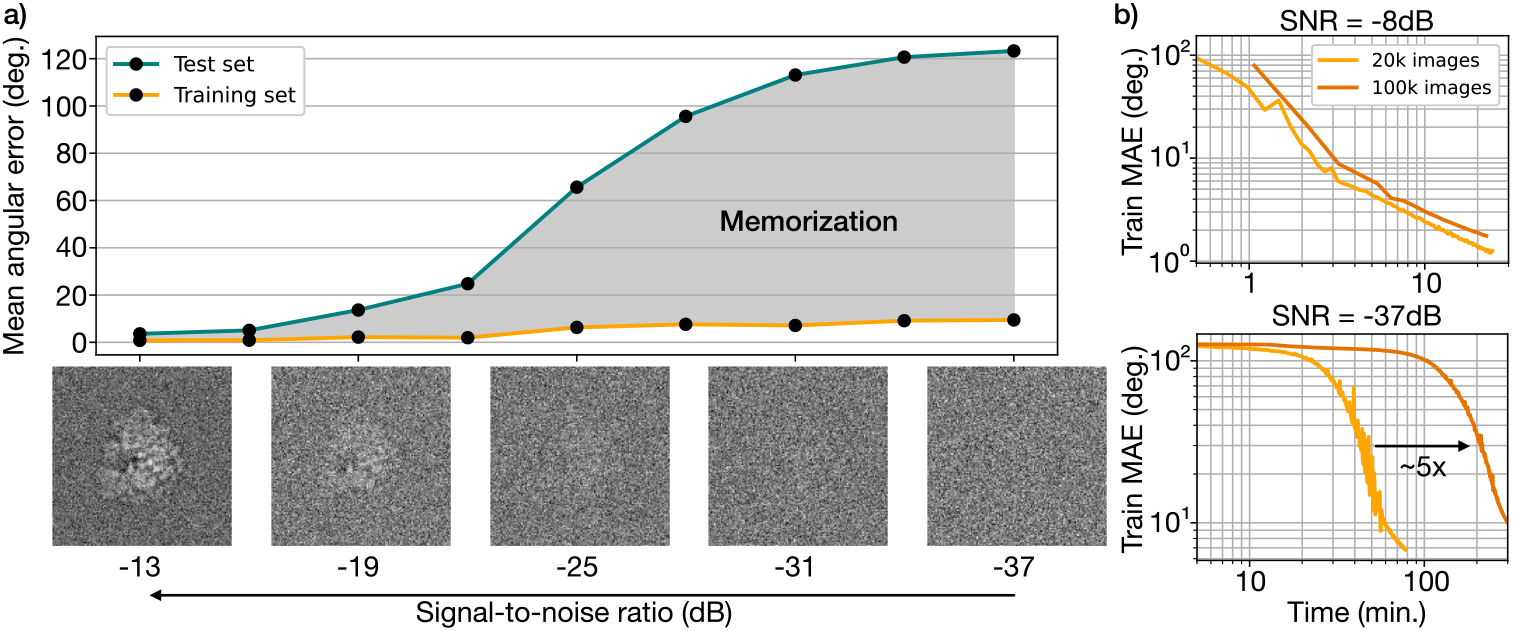
Memorization issue on high-noise datasets. **a)** Mean angular error of a neural-based pose predictor trained on increasingly noisy synthetic datasets. While the training-time error is stable, the gap with the test-time error increases with noise. **b)** Training-time mean angular error (MAE) as a function of time on small (20k images) and large (100k images) datasets. On low-noise datasets (*−*8 dB), convergence time does not depend on the dataset size, On high-noise datasets (*−*37 dB), convergence time increases with the dataset size. This “memorization cost” invalidates the benefits of the amortized approach on high-noise datasets. See Methods for experimental details.

**Supplementary Figure S2:**
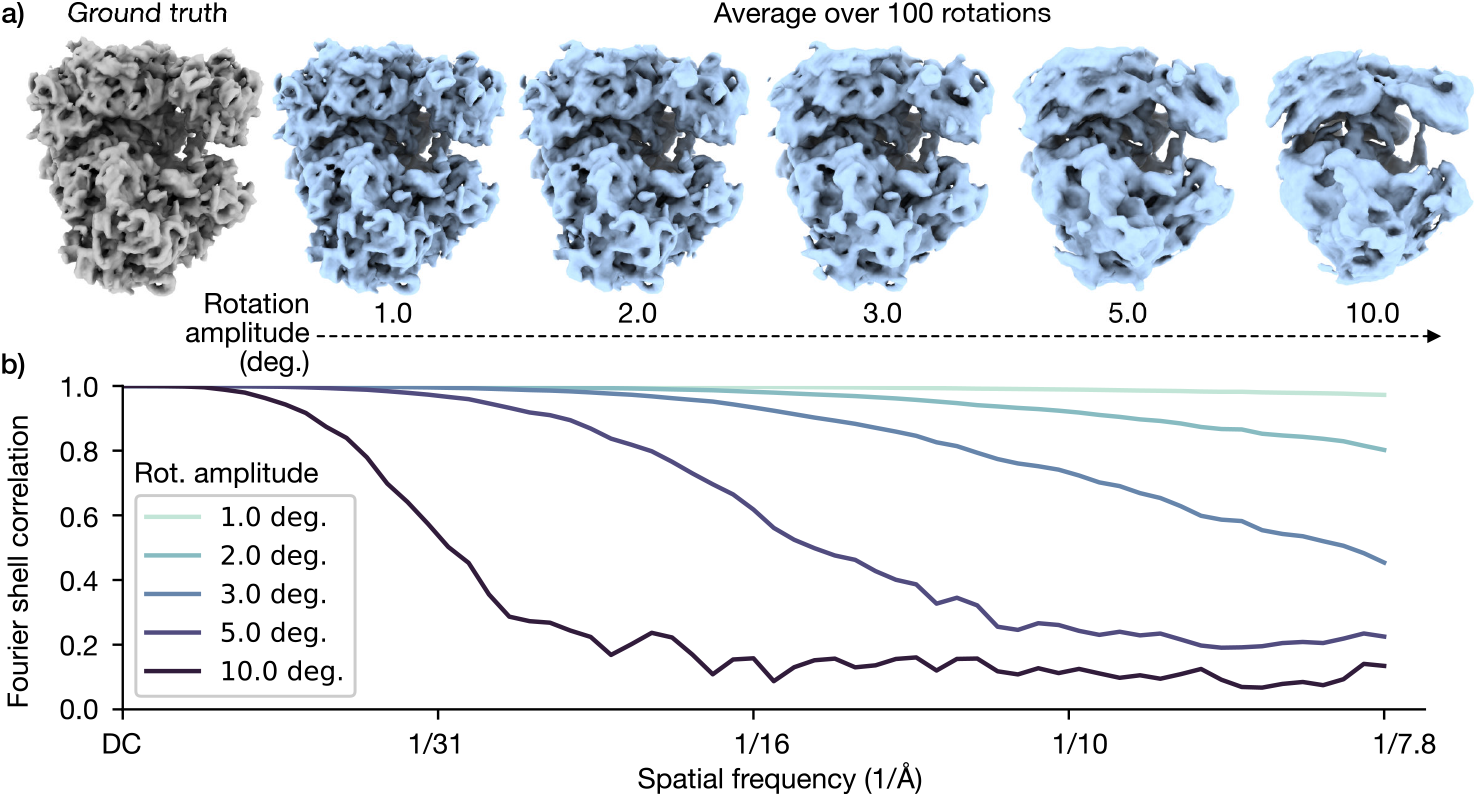
Influence of pose error on resolution. To illustrate the impact of inaccurate poses on the reconstructed density map, 100 randomly rotated density maps of the 80S ribosome (128×128×128, 3.77 Å/pix.) are averaged, using a rotation amplitude between 1 and 10 degrees. **a)** Ground truth and averaged density maps for different rotation amplitudes. **b)** Fourier shell correlation for different rotation amplitudes.

**Supplementary Figure S3:**
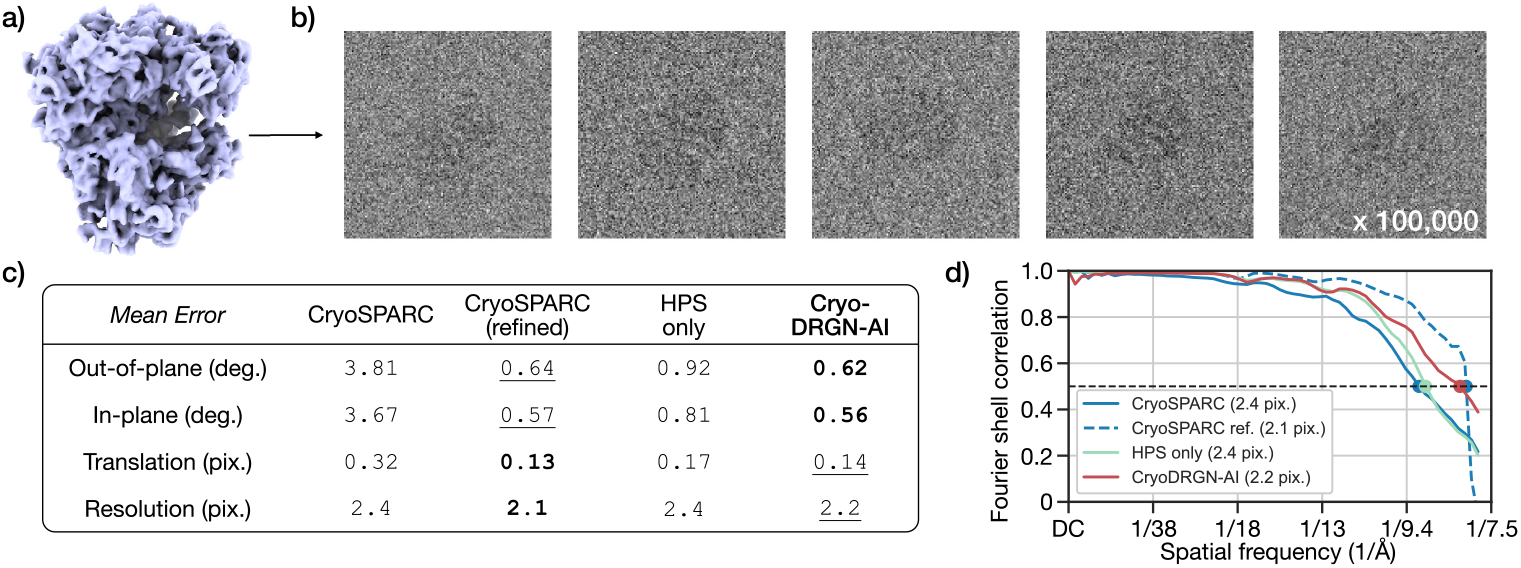
Homogeneous reconstruction of synthetic datasets. **a)** Ground truth density map of the 80S ribosome. **b)** Five example particle images (100,000 particles, 128×128, 3.77 Å/pix). **c)** Pose error for cryoDRGN-AI, cryoDRGN-AI using HPS only and cryoSPARC [15]. **d)** FSC curves between the reconstructed density map and the ground truth density map (resolution at FSC 0.5 between parenthesis).

**Supplementary Figure S4:**
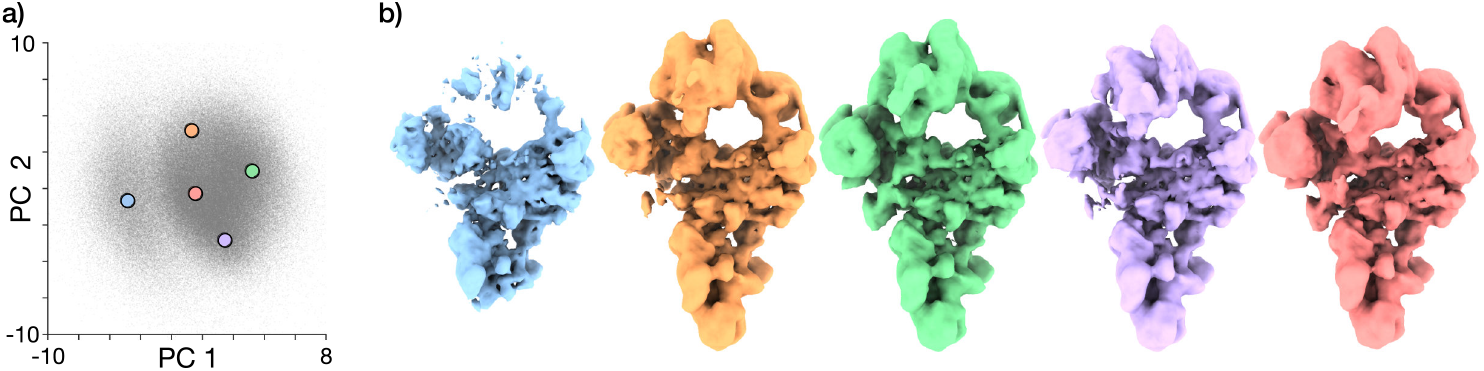
Additional visualizations of the latent space for the spliceosome dataset. **a)** UMAP visualization of the latent embeddings with 5 centroids obtained by k-means clustering. **b)** Associated density maps.

**Supplementary Figure S5:**
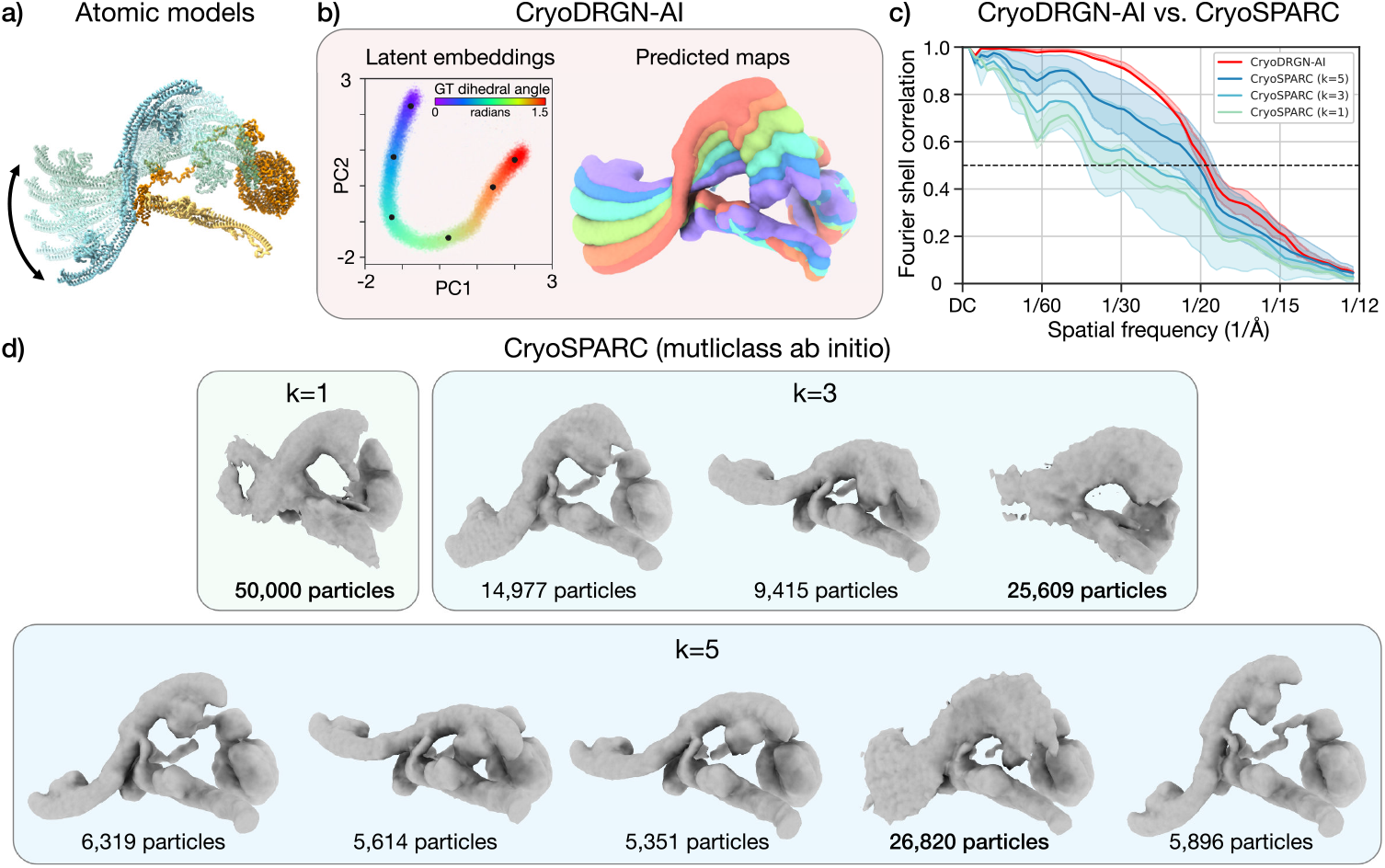
Results of cryoDRGN-AI on a synthetic dataset with strong conformational heterogeneity and comparison to cryoSPARC (multiclass *ab initio*). **a)** The dataset simulates strong conformational heterogeneity (see Methods). We show 5 models separated by a 0.3 rad. increment rotation along a dihedral angle. **b)** Output of cryoDRGN-AI. PCA on the latent embeddings on the left (the hue represents, for each image, the true dihedral angle) and associated density maps on the right (sampled on black dots). **c)** Comparison between cryoDRGN-AI and cryoSPARC multiclass *ab initio*. We show the mean and inter-quartile range of the per-image FSC computed on 50 images uniformly sampling the range of motion. **d)** Results obtained with cryoSPARC multiclass *ab initio* (default parameters), using different numbers of classes. We indicate the number of particles associated to each class and bold the highest number.

**Supplementary Figure S6:**
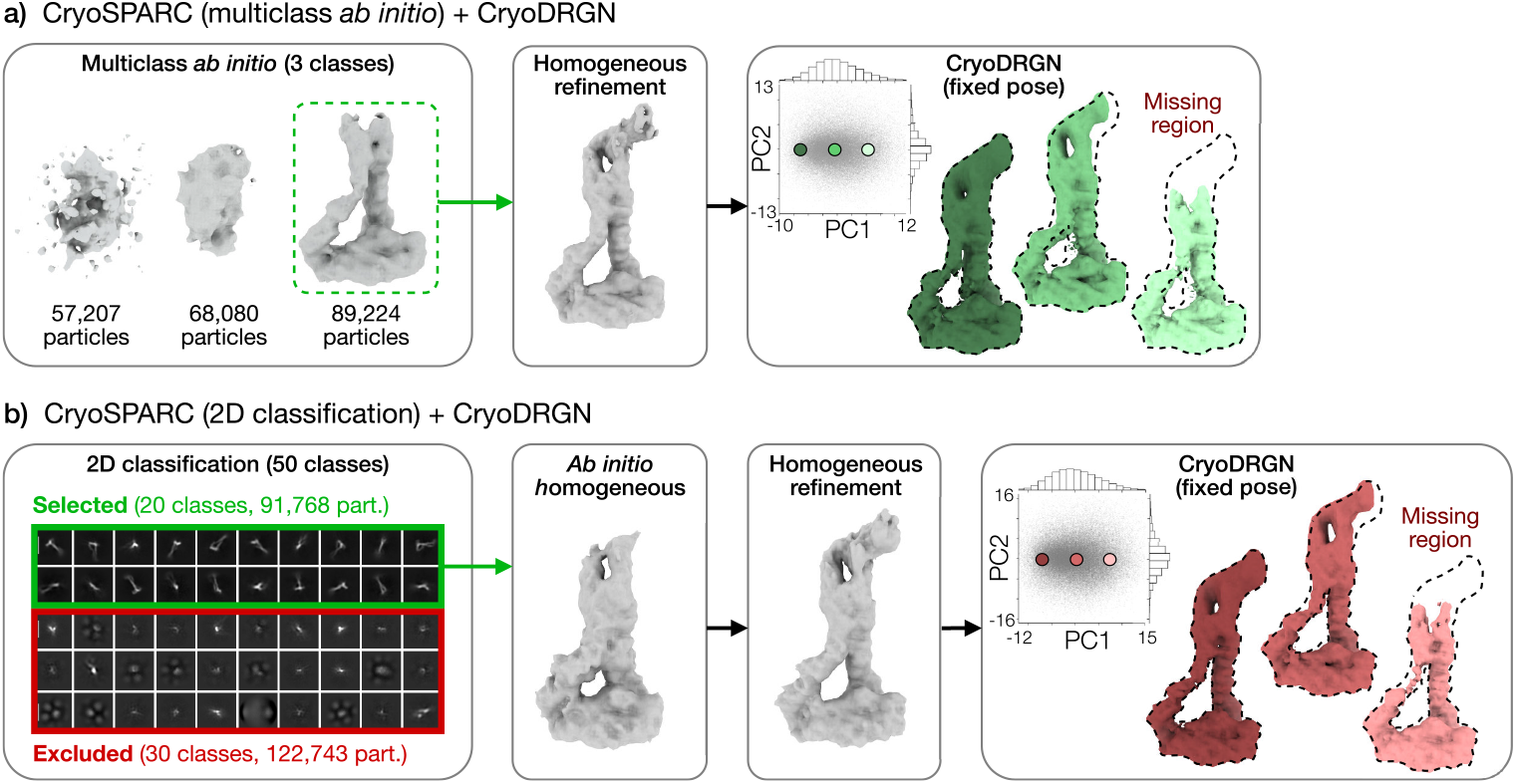
Comparison to other cryo-EM workflows for the DSL1 dataset, based on the cryoSPARC software [15]. **a)** Spurious particles are filtered using multiclass ab initio reconstruction with 3 classes. Poses are then refined with a step of homogeneous refinement. The filtered and posed particle stack is finally processed with cryoDRGN [5]. **b)** Filtering is done by 2D classification and followed by a step of ab initio reconstruction and a step of homogeneous refinement. The posed particle stack is then processed with cryoDRGN.

**Supplementary Figure S7:**
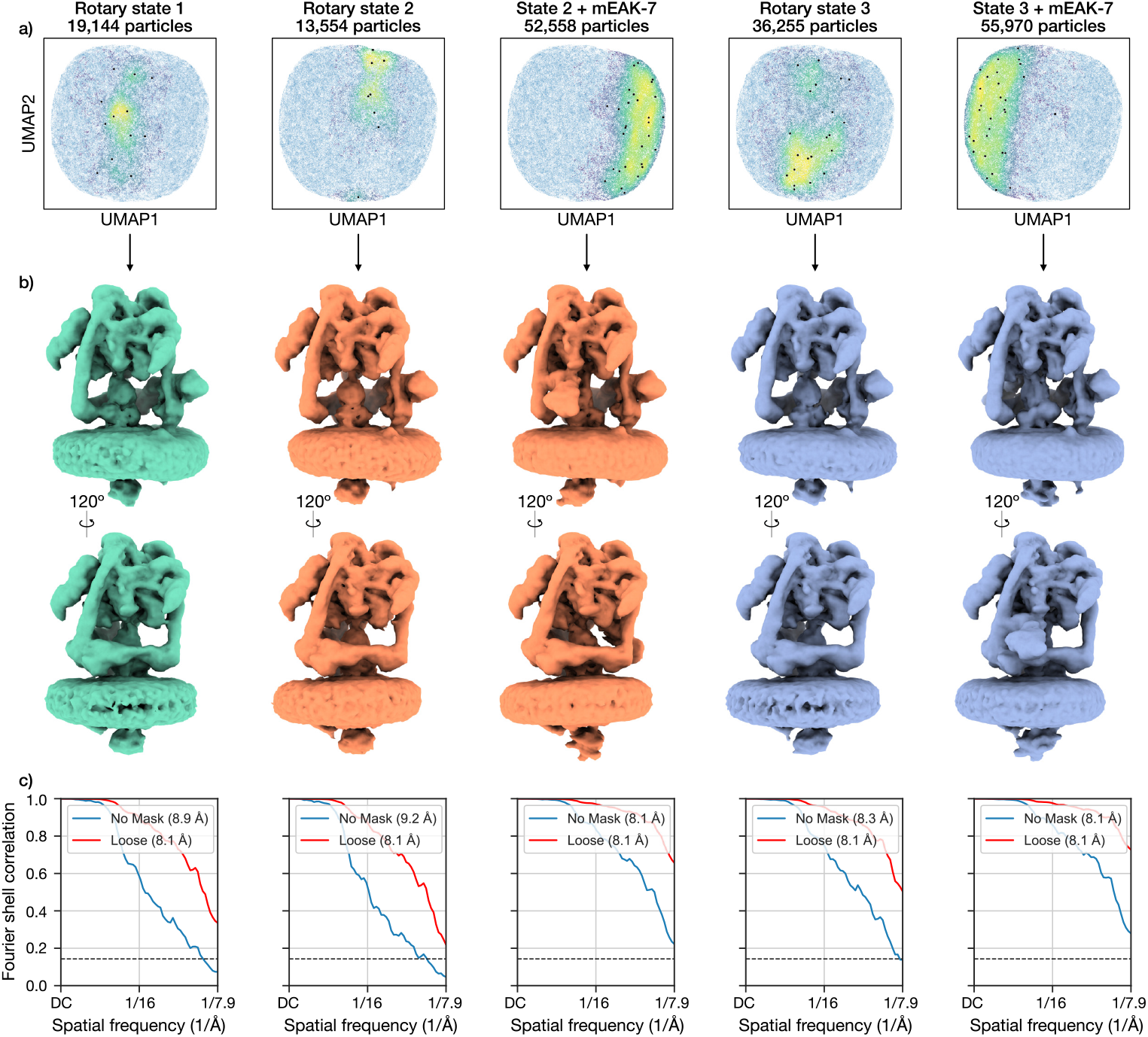
Backprojections of V-ATPase states. **a)** UMAP visualizations of the latent embeddings, colored by the Gaussian kernel density estimate of the selected particles from each state. **b)** Two views of backprojected density maps from the selected particles of each state. **c)** Half-map FSC with soft mask at half of maximum value, 15 Åcosine edge width, and 25 Ådilation.

**Supplementary Figure S8:**
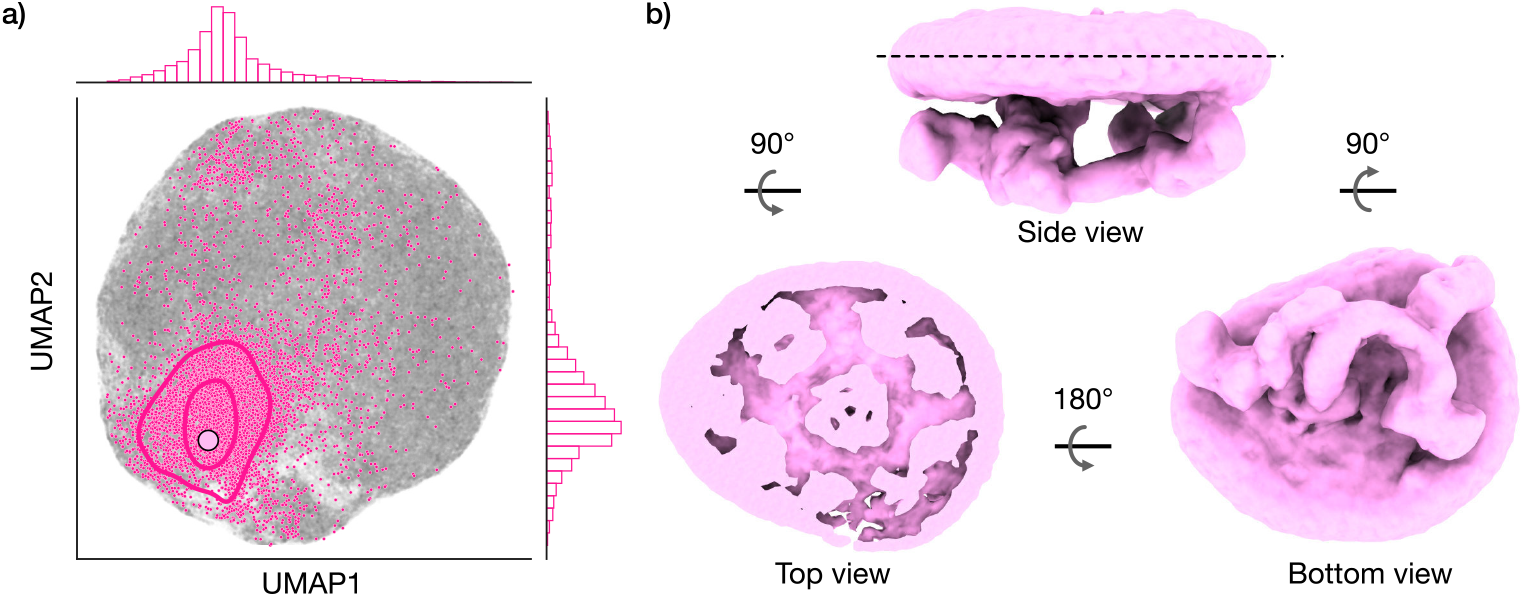
3D classification with cryoSPARC [15] on the ankyrin dataset, using the poses estimated by cryoDRGN-AI and 80 classes. **a)** UMAP of the latent embeddings with cryoDRGN-AI (grey). The large pink circle indicates the conformation of the supercomplex shown in Figure 5. The smaller dots indicate the conformations of the particles belonging to the supercomplex class, according to cryoSPARC (8,523 particles). These conformations are approximated with a Gaussian KDE shown with pink contours. **b)** Three views of the supercomplex state reconstructed with cryoSPARC “3D classification”, using cryoDRGN-AI-predicted poses (one class out of 80).

**Supplementary Figure S9:**
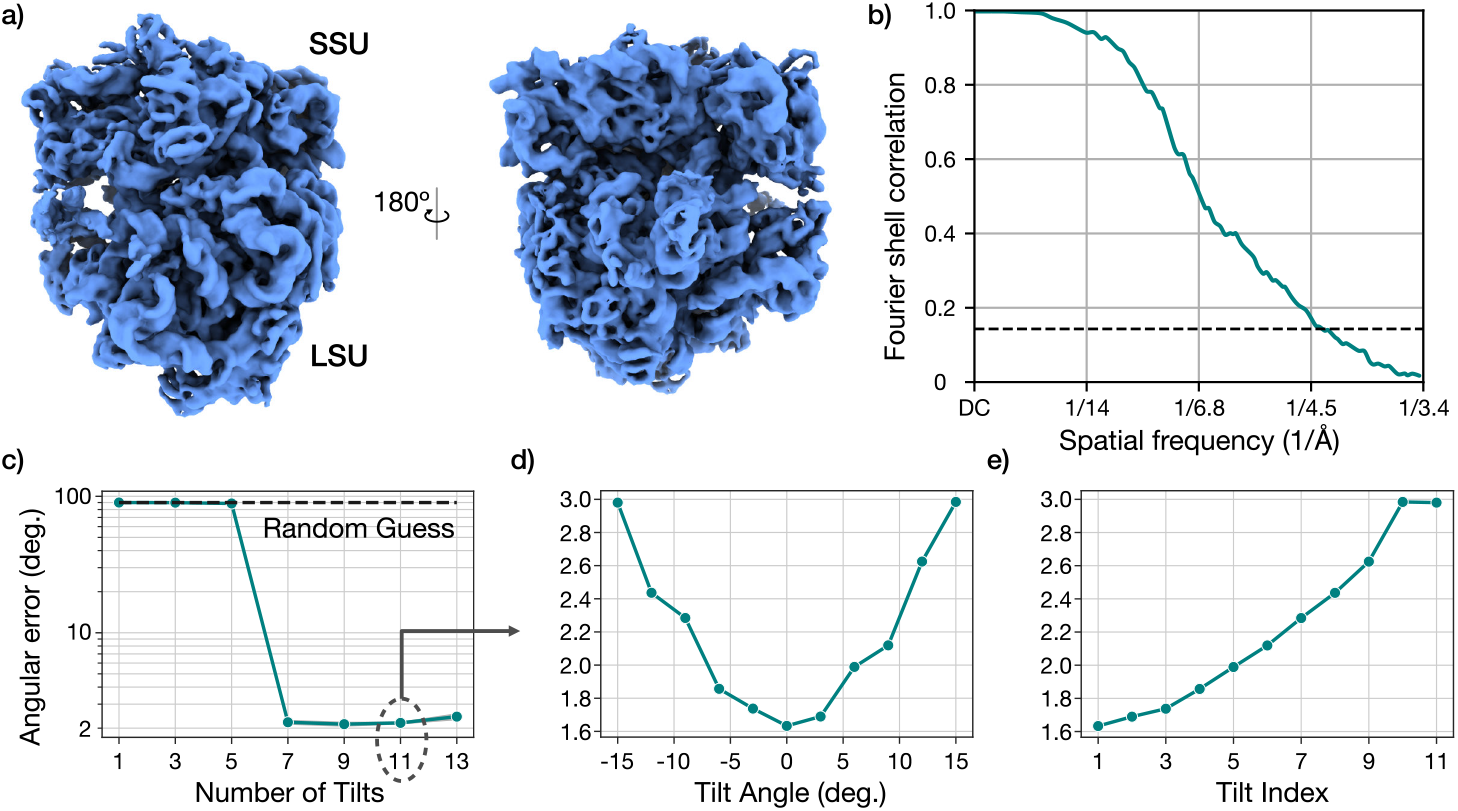
Homogeneous reconstruction of the “A, P stat” and pose accuracy of subtomogram reconstruction on the mycoplasma pneumoniae 70S ribosome. **a)** Reconstructed density map from 14,857 selected particles from Rangan et al. [12] (11 tilts per particle, 163,427 subtilt images total, 294×294, 1.70 Å/pix.). **b)** Half-map FSC with soft mask at half of maximum value, 15 Åcosine edge width, and 25 Ådilation. **c, d, e)** Mean out-of-plane angular error vs. number of tilts used for reconstruction (c), vs. tilt angle (d) and vs. tilt index (e). d and e were obtained using 11 tilts per particle.

**Supplementary Figure S10:**
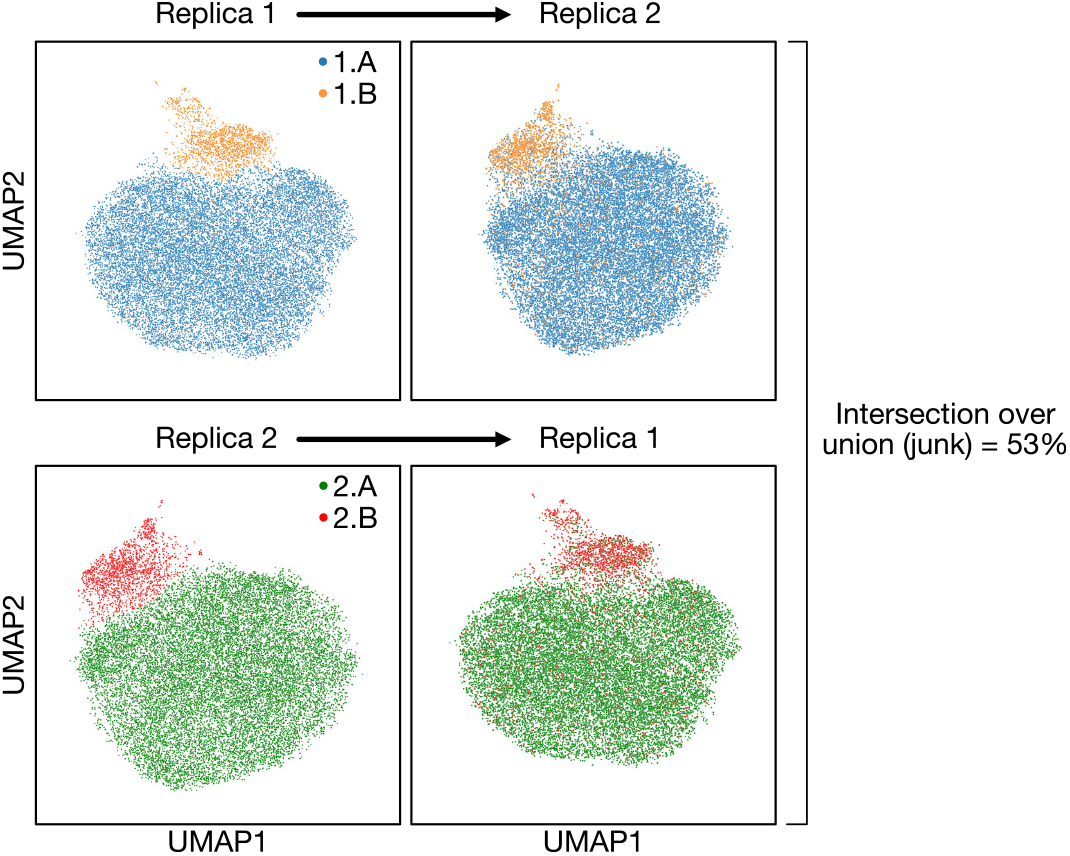
Consistent classification of junk in the mycoplasma pneumoniae 70S ribosome dataset. We run two independent experiments (replica 1 and 2) and manually select an outlying cluster in the UMAP plots (1.B and 2.B). Each particle is labeled according to the cluster it appears in after reconstruction 1 (resp. 2), and their distributions are shown in the UMAP plot of reconstruction 2 (resp. 1). Intersection over union would be 4.8% if the two classifications were uncorrelated.

## Data Availability

The following datasets can be accessed on EMPIAR (https://www.ebi.ac.uk/empiar/): pre-catalytic spliceo-some (EMPIAR-10180), assembling 50S ribosome (EMPIAR-10076), DLS1/SNARE complex (EMPIAR-11846), V-ATPase complex (EMPIAR-10874), ankyrin-1 complex (EMPIAR-11043), *Mycoplasma pneumoniae* 70S ribosome (EMPIAR-10499). Preprocessed particle stacks and synthetic datasets used in this work are deposited on Zenodo at https://doi.org/10.5281/zenodo.14853184 [80] (DLS1/SNARE complex), https://doi.org/10.5281/zenodo.14853225 [81] (V-ATPase complex), https://doi.org/10.5281/zenodo.14853246 [82] (*Mycoplasma pneumoniae* 70S ribosome), https://doi.org/10.5281/zenodo.14853257 [83] (synthetic 1D motion), https://doi.org/10.5281/zenodo.14853270 [84] (synthetic 80S ribosome). The covid spike dataset was provided by courtesy of the authors [18]. CryoDRGN-AI’s outputs were deposited on Zenodo at https://doi.org/10.5281/zenodo.14847271 [85]. Atomic models used from previous studies were obtained from the PDB (3J7A, 3J79, 6VXX, 6VYB, 8EKI).

## Code Availability

CryoDRGN-AI is available as open-source software at https://cryodrgnai.cs.princeton.edu and is integrated in version 4.0.0 of the cryoDRGN software package [86].

## References

[1] Nakane, T. et al. Single-particle cryo-em at atomic resolution. Nature 587, 152–156 (2020).

[2] Yip, K. M., Fischer, N., Paknia, E., Chari, A. & Stark, H. Atomic-resolution protein structure determination by cryo-em. Nature 587, 157–161 (2020).

[3] Frank, J. & Ourmazd, A. Continuous changes in structure mapped by manifold embedding of single-particle data in cryo-em. Methods 100, 61–67 (2016).

[4] Punjani, A. & Fleet, D. J. 3d variability analysis: resolving continuous flexibility and discrete heterogeneity from single particle cryo-em. J. Struct. Biol. 213, 107702 (2021).

[5] Zhong, E. D., Bepler, T., Berger, B., & Davis, J. H. Cryodrgn: Reconstruction of heterogeneous cryo-em struc-tures using neural networks. Nat. Methods 18, 176–185 (2021).

[6] Chen, M. & Ludtke, S. J. Deep learning-based mixed-dimensional gaussian mixture model for characterizing variability in cryo-em. Nat. Methods 18, 930–936 (2021).

[7] Nakane, T., Kimanius, D., Lindahl, E. & Scheres, S. H. Characterisation of molecular motions in cryo-em single-particle data by multi-body refinement in relion. eLife 7, e36861 (2018).

[8] Chua, E. Y. D. et al. Better, faster, cheaper: recent advances in cryo-electron microscopy. Annual Review of Biochemistry 91, 1–32 (2022).

[9] Oikonomou, C. M. & Jensen, G. J. Cellular electron cryotomography: toward structural biology in situ. Annual Review of Biochemistry 86, 873–896 (2017).

[10] Galaz-Montoya, J. G. & Ludtke, S. J. The advent of structural biology in situ by single particle cryo-electron tomography. Biophysics Reports 3, 17–35 (2017).

[11] Xue, L. et al. Visualizing translation dynamics at atomic detail inside a bacterial cell. Nature 610, 205–211 (2022).

[12] Rangan, R. et al. Cryodrgn-et: deep reconstructing generative networks for visualizing dynamic biomolecules inside cells. Nat. Methods 21, 1537–1545 (2024).

[13] Punjani, A. & Fleet, D. J. 3dflex: determining structure and motion of flexible proteins from cryo-em. Nat. Methods 20, 1–11 (2023).

[14] Zhong, E. D., Bepler, T., Davis, J. H. & Berger, B. Reconstructing continuous distributions of 3d protein struc-ture from cryo-EM images. In International Conference on Learning Representations (ICLR, 2020).

[15] Punjani, A., Rubinstein, J. L., Fleet, D. J. & Brubaker, M. A. Cryosparc: algorithms for rapid unsupervised cryo-em structure determination. Nat. Methods 14, 290–296 (2017).

[16] Davis, J. H. et al. Modular assembly of the bacterial large ribosomal subunit. Cell 167, 1610–1622 (2016).

[17] Plaschka, C., Lin, P.-C. & Nagai, K. Structure of a pre-catalytic spliceosome. Nature 546, 617–621 (2017).

[18] Walls, A. C. et al. Structure, function, and antigenicity of the sars-cov-2 spike glycoprotein. Cell 181, 281–292 (2020).

[19] DAmico, K. A. et al. Structure of a membrane tethering complex incorporating multiple snares. Nature Struc-tural & Molecular Biology 31, 246–254 (2024).

[20] Tan, Y. Z. et al. Cryoem of endogenous mammalian v-atpase interacting with the tldc protein meak-7. Life Science Alliance 5, (2022).

[21] Vallese, F. et al. Architecture of the human erythrocyte ankyrin-1 complex. Nature Structural & Molecular Biology 29, 706–718 (2022).

[22] Tegunov, D., Xue, L., Dienemann, C., Cramer, P. and Mahamid, J. Multi-particle cryo-em refinement with m visualizes ribosome-antibiotic complex at 3.5 Å in cells. Nat. Methods 18, 186–193 (2021).

[23] Bojanowski, P., Joulin, A., Lopez-Paz, D. & Szlam, A. Optimizing the latent space of generative networks. Preprint at https://arxiv.org/abs/1707.05776 (2017).

[24] Edelberg, D. G. & Lederman, R. R. Using vaes to learn latent variables: observations on applications in cryo-em. Preprint at https://arxiv.org/abs/2303.07487 (2023).

[25] Luo, Z., Ni, F., Wang, Q. & Ma, J. Opus-dsd: deep structural disentanglement for cryo-em single-particle analysis. Nat. Methods 20, 1729–1738 (2023).

[26] Gilles, M. A. T. & Singer, A. A bayesian framework for cryo-em heterogeneity analysis using regularized covariance estimation. Preprint at bioRxiv 10.1101/2023.10.28.564422 (2023).

[27] Zhong, E. D., Lerer, A., Davis, J. H. & Berger, B. Cryodrgn2: ab initio neural reconstruction of 3d protein structures from real cryo-em images. In Proceedings of the IEEE/CVF International Conference on Computer Vision 4066–4075 (CVPR, 2021).

[28] Herreros, D. et al. Estimating conformational landscapes from cryo-em particles by 3d zernike polynomials. Nature Communications 14, 154 (2023).

[29] Kaur., S. et al. Local computational methods to improve the interpretability and analysis of cryo-em maps. Nature Communications 12, 1240 (2021).

[30] McInnes, L., Healy, J. & Melville, J. Umap: Uniform manifold approximation and projection for dimension reduction. Preprint at https://arxiv.org/abs/1802.03426 (2018).

[31] Dashti, A. et al. Retrieving functional pathways of biomolecules from single-particle snapshots. Nature Com-munications 11, 4734 (2020).

[32] Maji, A. et al. Propagation of conformational coordinates across angular space in mapping the continuum of states from cryo-em data by manifold embedding. Journal of Chemical Information and Modeling 60, 2484–2491 (2020).

[33] Moscovich, A., Halevi, A., Andén, J. & Singer, A. Cryo-em reconstruction of continuous heterogeneity by laplacian spectral volumes. Inverse Probl. 36, 024003 (2020).

[34] Lederman, R. R. & Singer, A. Continuously heterogeneous hyper-objects in cryo-em and 3-d movies of many temporal dimensions. Preprint at https://arxiv.org/abs/1704.02899 (2017).

[35] Gupta, H., Phan, T. H., Yoo, J. & Unser, M. Multi-cryogan: reconstruction of continuous conformations in cryo-em using generative adversarial networks. In European Conference on Computer Vision (ECCV, 2020).

[36] Zhong, E. D., Lerer, A., Davis, J. H. & Berger, B. Exploring generative atomic models in cryo-em reconstruction. In NeurIPS Workshop on Machine Learning for Structural Biology (MLSB, 2020).

[37] Jin, Q. et al. Iterative elastic 3d-to-2d alignment method using normal modes for studying structural dynamics of large macromolecular complexes. Structure 22, 496–506 (2014).

[38] Harastani, M., Eltsov, M., Leforestier, A. & Jonic, S. Hemnma-3d: cryo electron tomography method based on normal mode analysis to study continuous conformational variability of macromolecular complexes. Frontiers in Molecular Biosciences 8, 663121 (2021).

[39] Hamitouche, I. & Jonic, S. Deephemnma: resnet-based hybrid analysis of continuous conformational hetero-geneity in cryo-em single particle images. Frontiers in Molecular Biosciences 9, 965645 (2022).

[40] Nashed, Y et al. Heterogeneous reconstruction of deformable atomic models in cryo-em. Preprint at https://arxiv.org/abs/2209.15121 (2022).

[41] Scheres, S. H. et al. Disentangling conformational states of macromolecules in 3d-em through likelihood opti-mization. Nat. Methods 4, 27–29 (2007).

[42] Elmlund, D. & Elmlund, H. Simple: software for ab initio reconstruction of heterogeneous single-particles. J. Struct. Biol. 180, 420–427 (2012).

[43] Scheres, S. H. Relion: implementation of a bayesian approach to cryo-em structure determination. J. Struct. Biol. 180, 519–530 (2012).

[44] Brubaker, M. A., Punjani, A., & Fleet, D. J. Building proteins in a day: efficient 3d molecular reconstruction. In Proceedings of the IEEE Conference on Computer Vision and Pattern Recognition 3099–3108 (CVPR, 2015).

[45] Ho, C.-M. et al. Bottom-up structural proteomics: cryoEM of protein com-plexes enriched from the cellular milieu. Nat. Methods 17, 79–85 (2020).

[46] Su, C.-C. et al. A “build and retrieve” methodology to simultaneously solve cryo-EM structures of membrane proteins. Nat. Methods 18, 69–75 (2021).

[47] Levy, A., Wetzstein, G., Martel, J. N., Poitevin, F. & Zhong, E. D. Amortized inference for heterogeneous reconstruction in cryo-em. In Advances in Neural Information Processing Systems (NeurIPS, 2022).

[48] Shekarforoush, S., Lindell, D. B., Brubaker, M. A. & Fleet D. J. Cryospin: improving ab-initio cryo-em recon-struction with semi-amortized pose inference. In Advances in Neural Information Processing Systems (NeurIPS, 2024).

[49] Tang, G. et al. Eman2: an extensible image processing suite for electron microscopy. J. Struct. Biol. 157, 38–46 (2007).

[50] Castaño-Díez, D., Kudryashev, M., Arheit, M. & Stahlberg, H. Dynamo: a flexible, user-friendly development tool for subtomogram averaging of cryo-em data in high-performance computing environments. J. Struct. Biol. 187, 139–151 (2012).

[51] Bharat, T. A. & Scheres, S. H. Resolving macromolecular structures from electron cryo-tomography data using subtomogram averaging in relion. Nature Protocols 11, 2054–2065 (2016).

[52] Harastani, M., Eltsov, M., Leforestier, A. & Jonic, S. Tomoflow: analysis of continuous conformational variabil-ity of macromolecules in cryogenic subtomograms based on 3d dense optical flow. J. Struct. Biol. 434, 167381 (2022).

[53] Himes, B. A. & Zhang, P. Emclarity: software for high-resolution cryo-electron tomography and subtomogram averaging. Nat. Methods 15, 955–961 (2018).

[54] Chen, M. et al. A complete data processing workflow for cryo-et and subtomogram averaging. Nat. Methods 16, 1161–1168 (2019).

[55] Tegunov, D. & Cramer P. Real-time cryo-electron microscopy data preprocessing with warp. Nat. Methods 16, 1146–1152 (2019).

[56] Powell, B. M. & Davis J. H. Learning structural heterogeneity from cryo-electron subtomograms with tomoD-RGN. Nat. Methods 21, 1525–1536 (2024).

[57] Zhu, J. et al. A minority of final stacks yields superior amplitude in single-particle cryo-EM. Nature Communi-cations 14, 7822 (2023).

[58] Kingma, D. P. & Welling M. Auto-encoding variational bayes. Preprint at https://arxiv.org/abs/1312.6114 (2013).

[59] Kinman L. F., Powell B. M., Zhong E. D., Berger B. & Davis J. H. Uncovering structural ensembles from single-particle cryo-EM data using cryoDRGN. Nat. Protocols 18, 319–339 (2023).

[60] Jeon, M. et al. Cryobench: diverse and challenging datasets for the heterogeneity problem in cryo-em. In Advances in Neural Information Processing Systems (NeurIPS, 2024).

[61] Jumper, J. et al. Highly accurate protein structure prediction with alphafold. Nature 596, 583–589 (2021).

[62] Abramson, J. et al. Accurate structure prediction of biomolecular interactions with alphafold 3. Nature 630, 493–500 (2024).

[63] Ingraham, J. B. et al. Illuminating protein space with a programmable generative model. Nature 623, 1070–1078 (2023).

[64] Watson, J. L. et al. De novo design of protein structure and function with rfdiffusion. Nature 620, 1089–1100 (2023).

## Methods-Only References

[65] Vulović, M. et al. Image formation modeling in cryo-electron microscopy. J. Struct. Biol. 183, 19–32 (2013).

[66] Tancik, M. et al. Fourier features let networks learn high frequency functions in low dimensional domains. In Advances in Neural Information Processing Systems (NeurIPS, 2020).

[67] He, K. Zhang, X., Ren, S. & Sun, J. Deep residual learning for image recognition. In Proceedings of the IEEE/CVF International Conference on Computer Vision 770–778 (CVPR, 2016).

[68] Paszke A. et al. Pytorch: an imperative style, high-performance deep learning library. In Advances in Neural Information Processing Systems (NeurIPS, 2019).

[69] Kingma, D. P. & Ba, J. Adam: A method for stochastic optimization. Preprint at https://arxiv.org/abs/1412.6980 (2014).

[70] Yershova, A., Jain, S., Lavalle, S. M. & Mitchell, J. C. Generating uniform incremental grids on so(3) using the hopf fibration. The International Journal of Robotics Research 29, 801–812 (2010).

[71] Gorski, K. M. et al. Healpix: a framework for high-resolution discretization and fast analysis of data distributed on the sphere. The Astrophysical Journal 622, 759 (2005).

[72] Grant, T & Grigorieff, N. Measuring the optimal exposure for single particle cryo-em using a 2.6 Å reconstruction of rotavirus vp6. eLife 4, e06980 (2015).

[73] Bharat, T. A., Russo, C. J., Löwe, J., Passmore, L. A & Scheres, S. H. Advances in single-particle electron cryomicroscopy structure determination applied to sub-tomogram averaging. Structure 23, 1743–1753 (2015).

[74] Pettersen, E. F. et al. Ucsf chimerax: structure visualization for researchers, educators, and developers. Protein Sci. 30, 70–82 (2021).

[75] Klindt, D. A., Hyvarinen, A., Levy, A., Miolane, N. & Poitevin F. Towards interpretable cryo-em: disentangling latent spaces of molecular conformations. Preprint at bioRxiv 10.1101/2024.03.18.585544 (2024).

[76] Zhou, Y., Barnes, C., Lu, J., Yang, J. & Li, H. On the continuity of rotation representations in neural networks. In Proceedings of the IEEE/CVF International Conference on Computer Vision 5745–5753 (CVPR, 2019).

[77] Levy, A. et al. CryoAI: amortized inference of poses for ab initio reconstruction of 3d molecular volumes from real cryo-em images. In European Conference on Computer Vision 540–557 (ECCV, 2022).

[78] Wong, W. et al. Cryo-em structure of the plasmodium falciparum 80s ribosome bound to the anti-protozoan drug emetine. eLife 3, e03080 (2014).

[79] Burt, A. et al. An image processing pipeline for electron cryo-tomography in relion-5. FEBS Open Bio 14, 1788–1804 (2024).

[80] Levy, A. et al. Input Data for dsl1/sanre complex in cryoDRGN-AI. Zenodo 10.5281/zenodo.14853184.

[81] Levy, A. et al. Input data for v-atpase complex in cryoDRGN-AI. Zenodo 10.5281/zenodo.14853225.

[82] Levy, A. et al. Input data for mycoplasma pneumoniae 70S ribosome in cryoDRGN-AI. Zenodo 10.5281/zenodo.14853246.

[83] Levy, A. et al. Input data for synthetic 1d motion in cryoDRGN-AI. Zenodo 10.5281/zenodo.14853257.

[84] Levy, A. et al. Input data for synthetic 80S ribosome in cryoDRGN-AI. Zenodo 10.5281/zenodo.14853270.

[85] Levy, A. et al. Output data for cryoDRGN-AI. Zenodo 10.5281/zenodo.14847271.

[86] Zhong, E. D. et al. ml-struct-bio/cryodrgn: v3.4.3 Zenodo 10.5281/zenodo.14538433.

